# Targeting *Pseudomonas aeruginosa* Ventilator-associated pneumonia with a non-antibiotic and biolfilm-disrupting Live Biotherapeutic: Preclinical Safety and Efficacy study

**DOI:** 10.1101/2025.08.24.671965

**Authors:** Laia Fernández-Barat, Ana Motos, Carolina Segura, Kasra Kiarostami, Roberto Cabrera, Alba Soler, Alicia Broto, Rocco Mazzolini, Flavia Galli, Blanca Llonch, Guillem Macip, Nil Vazquez, Davide Calabretta, Marta Arrieta, Maria Lluch, Montserrat Rigol, Luis Serrano, Irene Rodríguez-Arce, Antoni Torres

**Affiliations:** CELLEX research laboratories, CIBERES (Centro de Investigación Biomédica en Red de Enfermedades Respiratorias, 06/06/0028), Institut d’Investigacions Biomèdiques August Pi i Sunyer (IDIBAPS), Barcelona, Spain; University of Barcelona, Barcelona, Spain; Pulmonology Department, Hospital Clínic, Barcelona, Spain; Nantes Université, CHU Nantes, INSERM, Center for Research in Transplantation and Translational Immunology UMR 1064, Nantes, France; Centre for Genomic Regulation (CRG), The Barcelona Institute of Science and Technology, Dr. Aiguader 88, Barcelona 08003, Spain; Pulmobiotics Ltd, Barcelona, Spain; Universitat Pompeu Fabra (UPF), Barcelona, Spain; ICREA, Pg. Lluis Companys 23, Barcelona 08010, Spain

## Abstract

**Background.:** Ventilator-associated pneumonia (VAP) is the second most common hospital-acquired infection and is associated with high morbidity and mortality. The insertion of the endotracheal tube (ETT) compromises the host’s bronchial defenses and facilitates the formation of microbial biofilms and VAP. Although antibiotics remain the cornerstone treatment, their efficacy is often compromised by poor biofilm penetration and the prevalence of multidrug-resistant bacteria. This reveals the need to develop novel non-antibiotic therapeutic strategies. We engineered a non-pathogenic strain of the pulmonary bacteria *Mycoplasma pneumoniae* CV2 to express anti-biofilm and bactericidal enzymes (VAP platform). The therapeutic effect was validated in a murine model of *Pseudomonas aeruginosa* infection and *ex vivo* in ETTs of patients with *P. aeruginosa*-induced VAP.

**Methods.:** We evaluated the safety of an improved version of CV2 (CV8, Mycochassis) in a miniature pig model, and its efficacy with the VAP therapeutic platform (CV8_VAP) in a piglet model of *P. aeruginosa*-induced VAP. CV8_VAP was administered via intralobe instillation or nebulization. We assessed the platform’s ability to degrade biofilms formed on ETTs *in vivo* and its impact on the airway microbiota. We evaluated bacterial load in the lungs, inflammatory responses, histopathological and the clinical outcomes to provide a comprehensive characterization of therapeutic efficacy and safety.

**Findings.:** VAP platform reduced biofilm thickness and *P. aeruginosa* load in ETTs and lungs, with the nebulized route showing enhanced efficacy. CV8_VAP contributed on microbiome diversity restoration. Inflammatory markers in the lung were markedly decreased, suggesting a local immunomodulatory effect.

**Interpretation.:** These findings support that CV8_VAP as a promising, non-antibiotic therapeutic strategy for treating VAP caused by *P. aeruginosa*.

**Funding.:** The project that gave rise to these results has received funding from “La Caixa” Foundation (HR18-00058), and the European Research Council under the European Union’s Horizon 2020 research and innovation program (670216).

## Introduction

Ventilator-associated pneumonia (VAP) is the second most frequent cause of hospital-infection and carries high mortality and morbidity. Patients that develop VAP have higher mortality, higher length of mechanical ventilation, and Intensive Care Unit (ICU) and hospital stay compared to patients that not present this complication. The crude mortality of VAP ranges from 40 to 60%, but the attributable mortality is 13% (1). VAP pathogenesis involves multiple factors, including disruption of bronchial mechanical defenses by endotracheal tube (ETT) insertion, continuous micro aspiration of oropharyngeal and gastric contents, and early biofilm formation on ETTs, which perpetuates infection (2) .

The bacterial etiology of VAP depends on the duration of mechanical ventilation and patient-specific risk factors. Multidrug-resistant (MDR) Gram-negative bacteria—particularly *Pseudomonas aeruginosa*, carbapenem-resistant *Enterobacterales,* and *Acinetobacter* spp.—are responsible for up to 70% of cases in patients ventilated for more than 5 days or with additional risk factors (3–6). Among them, *P. aeruginosa* accounts for 20–30% of late-onset VAP and is the most frequent cause in many regions. Although antibiotics remain the cornerstone of VAP treatment (7), their effectiveness is increasingly limited by poor penetration into ETT-associated biofilms and the emergence of MDR strains (8). Additionally, a recent validation of current European international guidelines suggests that using empirically more than one antibiotic against Gram negative bacilli in patients with VAP and without shock, might increase mortality of these patients (9). Consequently, all measures to reduce the overuse and duration of antibiotics in VAP should be beneficial. This growing challenge underscores the urgent need for non-antibiotic therapies. This include monoclonal antibodies and live biotherapeutics (LBs), such as bacteriophages or engineered bacteria, designed to specifically target and eliminate resistant and non-resistant pathogens (10,11).

*Mycoplasma pneumoniae*, a causative pathogen of mild respiratory diseases, was repurposed to create a non-pathogenic chassis (CV2) for delivering therapeutic payloads. CV2 has been engineered to expresses anti-biofilm and bactericidal enzymes (VAP platform) with efficacy against *P. aeruginosa* biofilms *ex vivo* in ETTs of patients with VAP and in a murine lung infection model (12). Recently, we have improved this CV2 chassis by replacing the glyceraldehyde-3-phosphate dehydrogenase GlpD (*mpn051*; (13)) with GpsA, an enzyme with similar metabolic activity that produces H_2_O instead of H_2_O_2_, enhancing the therapeutic potential of the strain, referred to as CV8 or *Mycochassis* (14).

In this study, we first confirmed *Mycochassis* safety in miniature pigs. *Mycochassis* was then equipped with the VAP platform (CV8-Mycochassis), and its therapeutic potential was evaluated in mechanically ventilated piglets challenged with *P. aeruginosa*, using both bronchoscopy intralobe and nebulized administration routes (10). We investigated its effects across multiple domains, including microbiological parameters in ETTs and lung, histopathology, as well as clinical progression.

Our results demonstrated that CV8_VAP reduced biofilm thickness and *P. aeruginosa* load in ETTs, with minimal impact on the overall microbiome diversity. CV8_VAP also significantly reduced *P. aeruginosa* burden in both bronchoalveolar lavage fluid (BALF) and lung tissue, with the nebulized route showing enhanced efficacy.

Inflammation was markedly decreased in treated animals, suggesting a local immunomodulatory effect. Although lung histopathology scores remained unchanged, no adverse tissue effects were observed, supporting the safety of *Mycochassis*. These findings support CV8_VAP as a promising, non-antibiotic therapeutic strategy for treating VAP caused by *P. aeruginosa*.

## Material and methods

### Medium and strain growth conditions

For *M. pneumoniae* growth, Hayflick complete medium was prepared by mixing 800 ml of noncomplete medium A (20 g of PPLO broth (Difco), 30 g of Hepes (100 mM final)), 25 ml of 0.5% phenol red solution (Sigma-Aldrich), 200 ml of heat-inactivated horse serum (Life Technologies), 20 ml of sterile filtered 50% glucose and 1 ml of a 100 mg/ml stock of ampicillin (final concentration 100 µg/ml; Sigma-Alrich). Solid Hayflick 1% agar plates were prepared by mixing Bacto Agar (BD). When corresponding, gentamicin (Gm) (final concentration 100 µg/ml) or tetracycline (Tc) (final concentration 2 µg/ml) were added.

For *P. aeruginosa* growth, sterile Luria-Bertani broth was inoculated with a stationary phase culture of *P. aeruginosa* and incubated for 2 hours at 37°C to achieve 10^8^ CFU/mL in logarithmic growth phase. We used a derived strain from *P. aeruginosa* ATCC 2785, resistant to ceftriaxone (minimal inhibitory concentration, MIC>256 mg/L) and piperacillin-tazobactam (MIC >256 mg/L).

### Generation of CV8_VAP strain

The genes encoding the VAP payloads (12) were introduced into the chassis strain CV8 via sequential transformation using transposons. First, a transposon carrying the A1-II′, PelAh, and PslG genes was introduced, and a single clone was isolated. This clone was then transformed with a second transposon carrying the Pyocin L1 gene. Final construct was isolated by antibiotic selection (Gm, Tc). Crystal violet and growth curves against *P. aeruginosa* PAO1 and Boston 41501 strains were performed as described (12).

### Western Blotting

Mycoplasma strains grown to exponential phase in T-75 flasks were lysed using 1 mL of lysis buffer containing 4% SDS and 100 mM HEPES (Sigma-Aldrich). Lysates were sonicated on ice using a Bioruptor Plus (Diagenode) set to HIGH power for 10 minutes with 30-second ON/OFF cycles. Protein samples were separated by SDS– PAGE using Novex™ 4–12% Bis-Tris gels in MOPS running buffer (Thermo Fisher Scientific), followed by transfer to nitrocellulose membranes using the iBlot 3 Western Blot Transfer System (Thermo Fisher Scientific). Membranes were blocked with 5% skim milk in TBS-Tween and incubated with either a monoclonal anti-FLAG antibody (Sigma, F1804) to detect PelAh and PslAh, or custom-made anti-A1-II’ and anti-pyocin L1 polyclonal primary antibodies (Proteogenix). Detection was carried out using HRP-conjugated secondary antibodies (anti-rabbit IgG, Sigma A0545; or anti-mouse IgG, Sigma A9044). Signals were developed using the SuperSignal™ West Pico chemiluminescent substrate (Thermo Fisher Scientific) and visualized with the iBright CL750 Imaging System (Thermo Fisher Scientific).

### Ethics

The study was conducted at the University of Barcelona (UB) animal facilities, with protocol approval from the UB Animal Ethics Committee (159/20). All procedures complied with EU Directive 2010/63/UE, Spanish Real Decreto 53/2013, and adhered to PREPARE and ARRIVE guidelines. Experiments were carried out in Biohazard Level 2 facilities. Upon arrival, animals were housed (1–2 pigs/pen) and acclimatized for 4–7 days. Animal details are provided in **Supplementary Table 1**.

### Assessment of VAP strain safety

Four miniature pigs (Specipig SL; 30.8 ± 1.0 kg) were anesthetized with 2–2.5 mg/kg propofol and intubated with a 6.5 mm I.D. ETT (Ruschelit® Safety Clear Plus, Teleflex Inc., Limerick). Animals were ventilated using a SERVO-i ventilator (Maquet) in volume-control mode with initial settings: Volume Tidal (VT) 8 mL/kg, pressure trigger –2 cmH_2_O, FiO₂ 0.4, duty cycle 0.33, inspiratory rise time 5%, inspiratory pause 10%, positive end expiratory pressure (PEEP) 4 cmH_2_O, and respiratory rate (RR) adjusted for normocapnia. Inspired gases were conditioned via a heated humidifier (Fisher & Paykel) maintaining 37°C near the ‘Y’ piece; the inspiratory line was also heated. ETT cuff pressure was maintained at 28 cmH_2_O.

Sedation and analgesia were maintained with midazolam and fentanyl, with 2 mg/kg propofol boluses as needed. Femoral artery cannulation (ultrasound-guided) allowed for blood pressure monitoring and sampling; a Foley catheter (8–10 Fr.) was inserted for urinary drainage. Fluid therapy included Ringer’s lactate and 0.9% NaCl. Animals received ceftriaxone (1 g IV pre-intubation, then 75 mg/kg every 12 h), plus meropenem (25 mg/kg every 8 h) and vancomycin (15 mg/kg every 12 h) to prevent endogenous infection.

CV8 was cultured in T75 flasks with Hayflick medium for 3–4 days, washed, and disaggregated through a 25G syringe. A 50 mL bacterial suspension was prepared, and CFU counts confirmed via Hayflick agar plating. Minipigs were positioned in right lateral anti-Trendelenburg for inoculation. Using a flexible bronchoscope (Ambu® aScope™), 15 mL of suspension (10⁸ CFU) was instilled into right lung lobes (RCr, RM, RC) and 15 mL into left lobes (LCr, LC). Controls received PBS. Animals remained in right lateral position at 0° for 24 h to ensure right-sided colonization, then rotated every 6 h.

Every 6 h, hemodynamics and gas exchange were assessed. Blood pressures were measured using TrueWave Pressure Transducers (Edwards Lifesciences) leveled at mid-thorax. Tracheal aspirates were collected at 24, 48, and 72 h and cultured; further aspiration was based on clinical need. BAL was performed at 72 h. CBC, biochemistry, and coagulation tests were conducted every 12–24 h.

At 72 h, animals were euthanized with midazolam, fentanyl, propofol, and 60 mEq KCl. Lungs were excised in supine position, and two samples from affected lobes were collected. For histology, tissue was fixed in paraformaldehyde (24 h), ethanol (100%, 24 h), and stored in PBS at 4°C. For microbiology, lung tissue was homogenized and cultured using standard methods.

### Assessment of VAP strain efficacy

Twenty-five Large White-Landrace × Duroc pigs (36.38 ± 2.91 kg; MirRamadera, SL) were anesthetized with 2–2.5 mg/kg propofol and intubated with a 7.5 mm I.D. ETT (Ruschelit® Safety Clear Plus, Teleflex Inc.), following the procedures previously described. Animals were ventilated using a SERVO-i ventilator (Maquet) with initial settings: TV 8 ml/kg, PEEP 4 cmH_2_O, and RR adjusted for normocapnia.

Femoral artery and jugular vein cannulation enabled blood sampling and hemodynamic monitoring via an 8-Fr introducer and a 7-Fr Swan-Ganz catheter (Edwards Lifesciences). A 12-Fr Foley catheter was placed in the bladder. Antibiotic prophylaxis included ceftriaxone, vancomycin, and 100 mg/kg of piperacillin-tazobactam every 8 h.

Pigs were challenged with *P. aeruginosa* PAO1 (5 mL, ∼10⁸ CFU/mL) into the oropharynx via the subglottic aspiration system, immediately after surgery and again 4 h later. To prevent rapid aspiration, PEEP was set at 5 cmH_2_O and cuff pressure raised to 40 cmH_2_O during and 10 min after instillation. After 24 h, animals received *M. pneumoniae* CV8 or CV8_VAP via bronchoscopy (5 mL/lobe; total 25 mL) or nebulization (28 mL over 55– 75 min; 10⁸ CFU, accounting for ∼10% delivery loss).

Nebulization was performed with a vibrating mesh nebulizer (In-line eFlow, PARI) 15 cm from the Y-piece. Ventilator settings were adjusted: RR halved (≥14 breaths/min), inspiratory rise time set to 0%, pause increased to 20%, I:E ratio to 1:1, and PEEP to ≥5 cmH_2_O. Pressure trigger sensitivity was set to –10 cmH_2_O. Sedation and analgesia doses were increased by 20%. Nebulization duration was recorded, and arterial blood gases were analyzed before, 15 min after, and at completion. Adverse events (e.g., bronchospasm, desaturation) were monitored.

Every 6 h, gas exchange and hemodynamic parameters were assessed. Central and arterial pressures, pulmonary artery pressure, wedge pressure, and cardiac output (via thermodilution) were recorded. Stroke volume, vascular resistances, and venous admixture were calculated. Ventilator settings were adjusted based on blood gases. Crystalloid boluses were given if MAP <65 mmHg, and norepinephrine administered for persistent hypotension. CBC, biochemistry, and coagulation tests were performed every 12–24 h.

BALF was performed with 10 mL 0.9% NaCl per lobe. Fluid was centrifuged (15,000 rpm, 5 min, 4°C); supernatant stored at –80°C for cytokine assays, pellets for RNA or bacterial load. Lung tissue was collected for histology, CFU count, and microbiome profiling.

Pigs were euthanized if PaO₂/FiO₂ <70 mmHg, if septic shock was unresponsive to inotropes, or at study end (36 h post-treatment), via anesthetic overdose and 60 mEq KCl IV.

### ETT slicing

Each 7.5 mm I.D. ETT was processed under sterile conditions to generate four sample types for different analyses. The cuff was removed for quantitative microbiological culture of external contaminants. The outer ETT surface was then cleaned with sterile gauze and rinsed with 80% ethanol and 0.9% saline (Braun). Three cross-sectional samples were taken from distal to proximal ends. Two 1-cm sections were divided into 0.5-cm pieces for confocal laser scanning microscopy (CLSM) and scanning electron microscopy (SEM). An adjacent 3-cm section was used for quantitative culture and microbiome analysis of inner biofilm colonization (**Supplementary** Figure 1) (13).

### Scanning Electron Microscopy (SEM)

Biofilm was imaged and thickness measured via SEM at the Electron Microscopy Unit-CCiTUB of the Faculty of Medicine of the UB. Minimal and maximal biofilm thickness was measured using dedicated software (ImageJ, Wayne Rasband). Concerning sample processing, briefly, the1-cm-long hemi-sections of the ETT distal dependent parts were fixed into a 2.5% glutaraldehyde and 2% paraformaldehyde-buffered solution followed by osmium tetroxide (1%) and potassium ferricyanide (0.8%). Subsequently, samples were dehydrated in graded ethanol series, dried using a polaron critical point drying apparatus, and mounted on commercial SEM stubs (Ted Pella). To avoid charge artefacts, the sections were sputter-coated with a gold thin layer (sc 510, Fisons Instrument) and carefully silver painted. SEM images were analyzed to determine maximum, minimum and mean thickness (µm) across different treatment groups.

### Confocal Laser Scanning Microscopy (CLSM)

The 0.5-cm cross-sections of ETTs were immersed in a 1 mL PBS, stained with live/dead® BacLight kitTM (BacLight kitTM; Invitrogen) for 15 min protected from the light, and then rinsed with PBS. The staining conditions were as follows: 1.5 µL of SYTO® 9 (stock 3.34 mM DMSO) and 1.5 µL propidium iodide (stock 20 mM DMSO) in 1 mL PBS. During CLSM imaging, SYTO® 9 fluoresces green and is used to identify living microorganisms with intact membranes whereas propidium iodide (PI) fluoresces red and stains dead bacteria with a damaged membrane. A Leica TCS SP5 laser scanning confocal system (Leica) of the Advanced Optical Microscopy Unit-CCiTUB of the Faculty of Medicine of the UB was used, containing a DMI6000 inverted microscope and a 20xPL APO numerical aperture 0.7 objectives. SYTO® 9 and PI images were acquired sequentially using 488-561-nm laser lines, an acoustic-optical beam splitter and emission detection range 500– 550, 570–620 nm, respectively. The confocal pinhole is set at 1 Airy unit. The pixel size was 160 nm. All samples and slides were coded to ensure that the image acquisition and measurements were blinded. Images were edited through ImageJ (Wayne Rasband).

### DNA extraction from ETT and Lung biopsies

One mL of the same sample used for microbiological analysis (i.e. the ETT previously resuspended in physiological saline solution 0.9%) was used to extract DNA through Spin columns by Sputum DNA kit Isolation (Norgen Corp), according to the manufacturer’s instructions with slight modifications to optimize the extraction of DNA (22). DNA was subsequently quantified by Qubit 4 Fluorometer (Thermo Fisher Scientific) and stored at −80°C upon further analysis.

For microbiome analysis of lung samples, tissue fragments were collected from each lobe, with five fragments per lobe (N=30 fragments). The fragments were pooled and homogenized on ice in 5 ml of PBS using an Ultra-Turrax (IKA). One milliliter of the homogenate was collected and centrifuged (5 min, 10000 rpm, 4°C), and the resulting pellet was used for DNA extraction (DNeasy UltraClean Microbial Kit, Qiagen), following the manufacturer’s instructions. DNA concentrations were measured in a Nanodrop One (Thermo-Scientific), and DNA integrity was assessed via 1% agarose gel electrophoresis.

DNA libraries preparation was based on the hypervariable regions V3-V4 of bacterial 16S rRNA and performed using the QIASeq Screening Panel 16S. The obtained pool of DNA libraries was sequenced using the Illumina MiSeq Sequencing System at IDIBAPS Core facilities. The results of each sample were obtained in fastq format files to perform bioinformatic microbiome analyses (15).

Data was processed using nf-core/ampliseq version 2.11.0 of the nf-core collection of workflows(16), utilizing reproducible software environments from the Bioconda and Biocontainers projects (17,18). Data quality was evaluated with FastQC and summarized with MultiQC (19). Cutadapt (20)trimmed primers and all untrimmed sequences were discarded. Sequences that did not contain primer sequences were considered artifacts. Adapter and primer-free sequences were processed sample-wise (independent) with DADA2 (21)to eliminate PhiX contamination, trim reads (before median quality drops below 25 and at least 75% of reads are retained; forward reads at 276 bp and reverse reads at 274 bp, reads shorter than this were discarded), discard reads with > 2 expected errors, correct errors, merge read pairs, and remove PCR chimeras. The microbiome analysis of lung tissue presented additional challenges, necessitating further filtering. Genomic DNA sequences from pig origin were identified and excluded from the bioinformatics analysis to ensure accurate results.

Taxonomic classification was performed by DADA2 and the database ‘Silva 138.1 prokaryotic SSU’(21). ASV sequences, abundance and DADA2 taxonomic assignments were loaded into QIIME2 (29). Within QIIME2, the final microbial community data was visualized in a barplot, evaluated for sufficient sequencing depth with alpha rarefaction curves, investigated for alpha (within-sample) and beta (between-sample) diversity after rarefaction and used to find differential abundant taxa with ANCOM-BC. Additional statistical tests were performed in Rstudio (RStudio Team, 2024) using R (R Core Team, 2024), with the packages dplyr and Hmisc.

Alpha diversity was quantified by observed species, Shannon and inverse Simpson’s diversity indices, and beta diversity through the Bray-Curtis dissimilarity index. Finally, the total bacterial DNA, the relative frequencies of *P. aeruginosa* and *M. pneumoniae* as well as other autochthonous mycoplasma species from the pig (*Mycoplasma hyopneumoniae* and *Mycoplasma hyorhinis*) recovered from ETTs were reported.

### Quantitative microbiology analysis

ETTs were processed and analyzed between November 2021 and November 2023 immediately after animal autopsy. All 3-cm cross sections and cuffs were immersed in 3 mL physiological saline solution 0.9% and sonicated in ultrasonic cleaning equipment (Branson 3510 E-MT; Bransonic) for 5 min at 40 kHz. Each solution was serially diluted, plated on Columbia Agar with 5% Sheep Blood (Becton Dickinson GmbH), and MacConkey Agar II (Becton Dickinson GmbH) and incubated overnight in a 5% CO2 incubator (Thermofisher Waterjacketed) for 24 h at 37°C. Bacterial growth was quantified and reported as the Median [IQR] of the logarithmic scale of CFU/ml (Log10 CFU/mL). Furthermore, all observed morphologies were isolated and identified by MALDI-TOF (Matrix-Assisted Laser Desorption/Ionization - Time of Flight) Mass Spectrometry.

Quantitative bacterial cultures of tracheal aspirates, bronchoalveolar lavage, and lung tissue were performed using standard methods. Ultimately, bacteria were identified by mass spectrometry through a Microflex LT (BrukerDaltonics GmbH, Leipzig, Germany) and bacterial identification was performed using the MALDI BioTyper 2.0 software (BrukerDaltonics).

### Macroscopic and histopathological assessment

For the macroscopic analysis of lung tissue, images were taken from the dorsal and ventral sides of all studied lungs. The background was removed using Figma and the pictures were normalized for brightness. Pictures were then converted to grayscale using the Python PIL library, ensuring that they were represented in a single channel where pixel values ranged from 0 (black) to 255 (white). Next, the images were transformed into NumPy arrays, allowing for efficient and effective numerical operations on the pixel values. The average pixel value of the remaining grayscale tones was calculated for each side, taking into account the total number of pixels. Finally, the mean of the values from both the dorsal and ventral sides was calculated. All images were evaluated on the same bit scale.

Tissues were embedded in paraffin using (Tec2900 Histo-Line Workstation), sectioned into 4 μm slices with a Leica RM2235 Microtome, and stained with hematoxylin and eosin (H&E) following standard protocols using (Leica Autostainer XL). The stained sections were manually mounted on glass slides and analyzed by light microscopy. Blinded evaluations were conducted at the Histopathology Facility of the Institute for Research in Biomedicine (IRB) in Barcelona, applying a previously established quantitative scoring system ((10). For the efficacy study, samples were ranged from 1 to 5, where: 0 - No pneumonia, 1-Purulent mucous plugging, 2 – Bronchiolitis, 3-Pneumonia, 4 - Confluent pneumonia, and 5 - Abscessed pneumonia. For each animal, five lung samples were assessed, representing specific lung regions (RCr, RM, RC, LCr, LC), and the final score was calculated as the average of these assessments.

### Statistical analysis

Data were processed with IBM SPSS Statistics for Windows, version 26.0 (IBM Corporation) or GraphPad Prism version 9.1.1 (GraphPad Software). Statistical analysis performed is specified in each figure legend. In all cases, p-value ≤ 0.05 was considered statistically significant.

## Results

### Testing CV8 safety in minipigs

The attenuation of the CV8 chassis has been previously validated in mouse lung models (14). However mouse lungs and immune system are very different from that of humans and therefore dose, distribution and pathology need to be assessed in a system close to humans. Pigs closely resemble humans in respiratory tract size, anatomy, physiology, and immune responses, addressing the limitations of murine models.

We tested the safety of the CV8 strain in a minipig model. In brief, minipigs were intubated and administered 10^8^ CFUs of CV8 or PBS in both the right and left lung lobes. Pigs were clinically monitored and sacrificed at 72 h post-infection for evaluation of the tissue damage (Figure 1 and Supplementary Table 2). Although macroscopic differences may be observed between CV8 and control animals (Figure 1A), evaluation of lung tissue showed no pneumonia or other relevant histopathological findings, confirming the attenuation of the CV8 strain (14) (Figure 1B, Supplementary Table 2). Clinically, the animals remained stable throughout the procedure, with no significant differences observed in respiratory (Figure 1C-G) or hemodynamic parameters (Figure 1H-I) compared to the control group and overtime.

**Figure 1.**
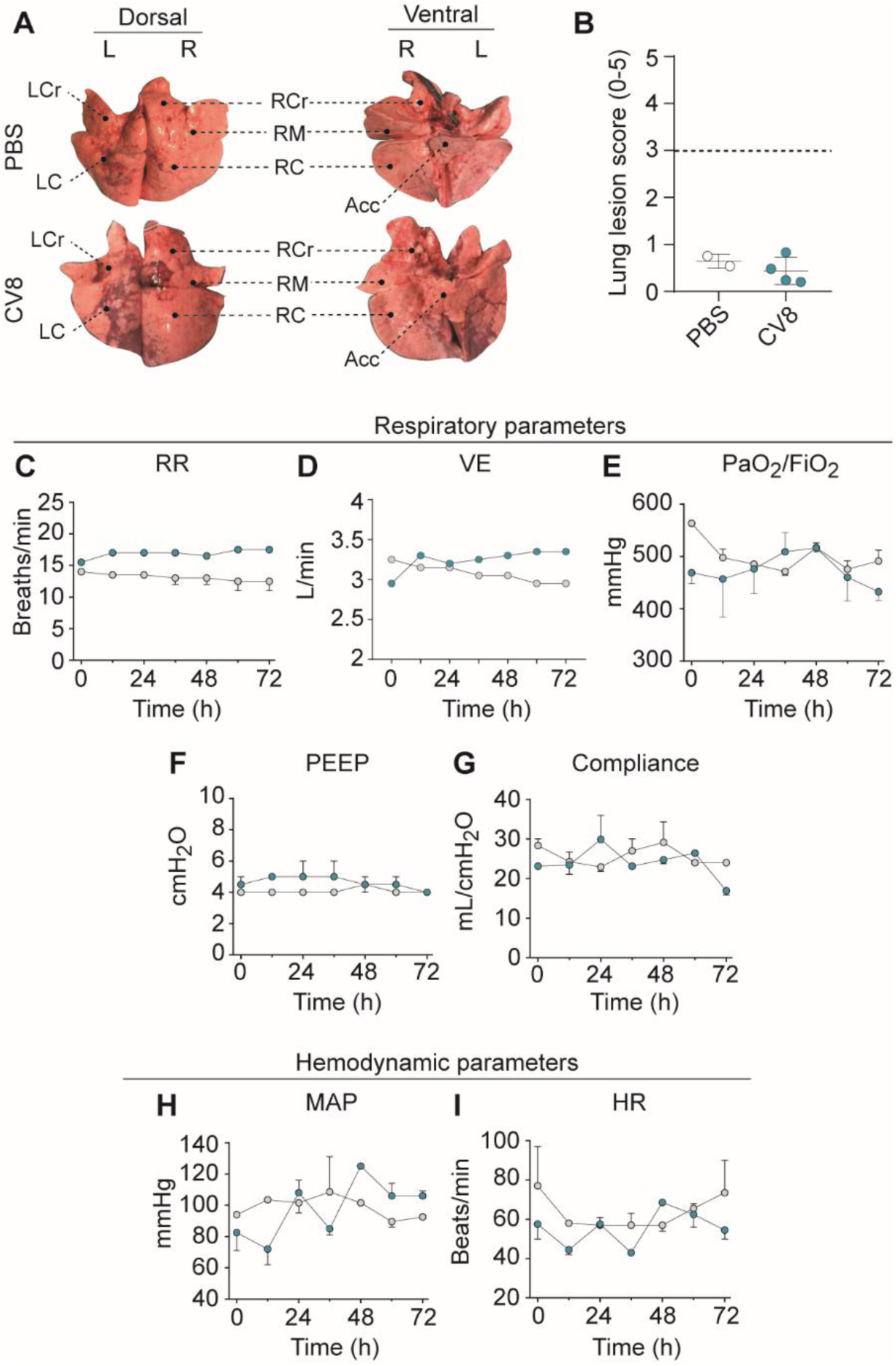
Safety assessment of the CV8 *Mycochassis* in minipigs. (**A**) Macroscopic images of right (R) or left (L) lungs from minipigs treated with PBS (top) or CV8 (bottom). Right lung cranial (RCr), middle (RM), caudal (RC); and left cranial (LCr) and caudal (LC) are shown. (**B**) Histological lesions observed in lung tissue treated with PBS (grey) or CV8 (blue). Each dot represents the average lesion score of lung lobes evaluated per animal (PBS, N=2; CV8, N=4) (see Supplementary Table 2). The dotted line (score = 3) indicates pneumonia. Respiratory (**C–G**) and hemodynamic (**H-I**) parameters assessed over 72 h in minipigs inoculated with PBS (grey) or CV8 (blue). (**C**) Respiratory rate (RR, breaths/min), (**D**) Volume minute (VE, L/ min), (**E**) PaO₂/FiO_2_ levels (mmHg), (**F**) Positive End-Expiratory Pressure (PEEP, cmH_2_O), (**G**) Respiratory compliance (mL/cmH_2_O), (**H**) Mean of Arterial Pressure (MAP, mmHg) and (**I**) Heart rate (HR, beats/min) are shown. Data are presented as Mean ± SEM.

### Effect of VAP platform treatment in ETTs biofilm architecture

After confirming its attenuation, we engineered CV8 biofilm-dispersing and antimicrobial functions by inserting the VAP platform (CV8_VAP), previously validated *in vivo* in a mouse model of *P. aeruginosa* PAO1 infection (Supplementary Figure 2) (12).

We evaluated the effect of CV8_VAP on *P. aeruginosa* PAO1 biofilms formed *in vivo* when administered as a treatment in a porcine model of VAP.

First, we conducted a quantitative measurement of biofilm thickness in the ETT from SEM images. The control group PBS + PBS showed the highest minimum, maximum, and mean biofilm thickness compared to the PAO1-infected groups (Figure 2A, Table 1). This biofilm is expected to consist of commensal bacteria from the oropharynx that can colonize the ETT. In PAO1-infected groups, CV8_VAP treatment tended to result in lower, although non-significant, values compared with PAO1 + PBS or CV8, particularly for maximal and mean thickness. This trend may be related to the effect of the therapeutic platform (Figure 2A, Table 1). Interestingly, minimal biofilm thickness appeared to reveal the largest differences among groups. The PAO1 + PBS group showed the highest value, whereas CV8 and CV8_VAP treatments were associated with a significant reduction (Figure 2A, Table 1).

**Figure 2.**
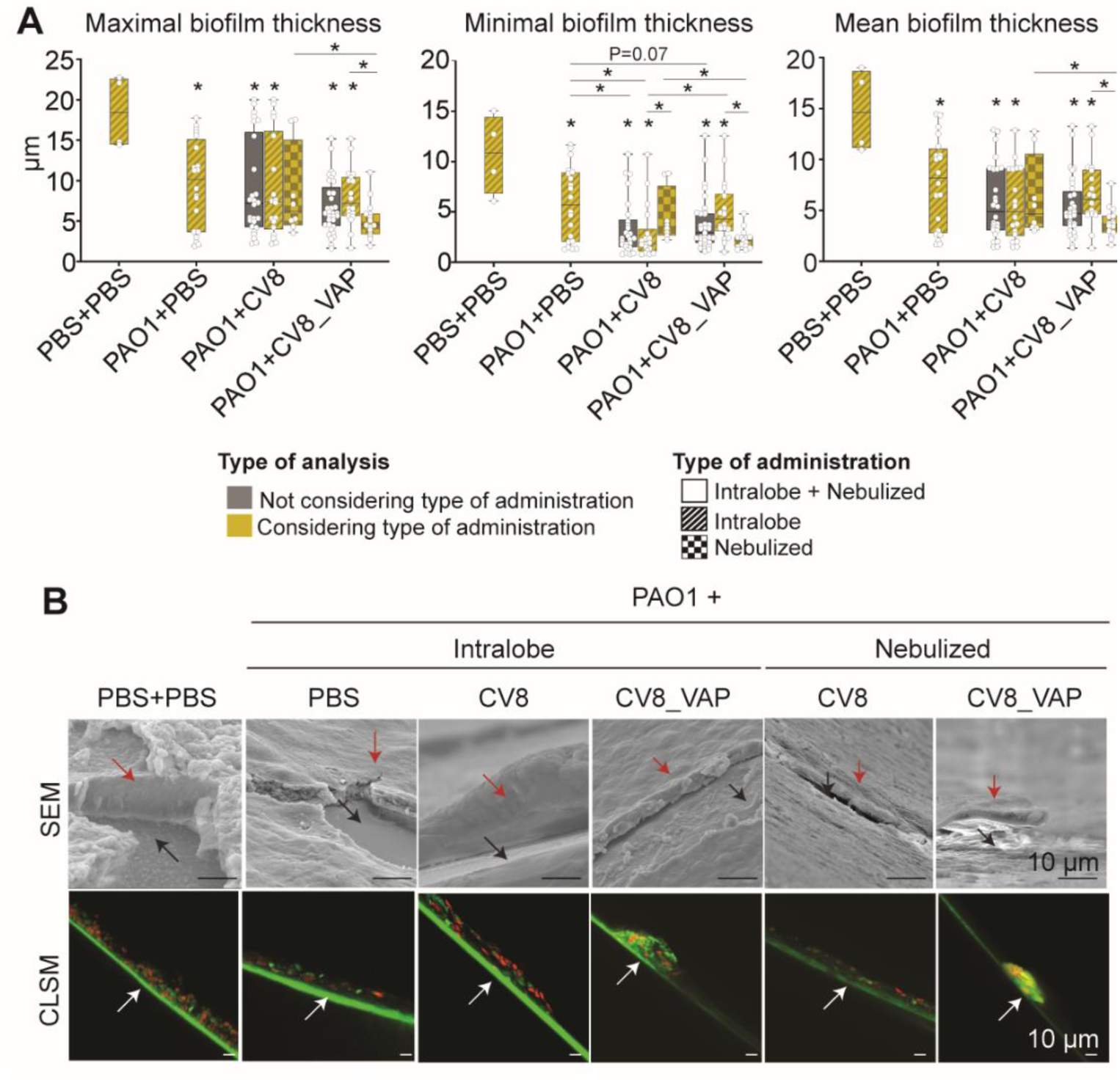
*In vivo*–formed ETT biofilm thickness after treatment with PBS, CV8 or CV8_VAP. **(A)** Maximal (left), minimal (middle), and mean (right) biofilm thickness determined by SEM. Experimental groups and number of images analyzed were as follows: PBS + PBS (n = 4; intralobe), PAO1 + PBS (n = 20; intralobe), PAO1 + CV8 (n = 26; intralobe, n = 18, nebulized, n = 8), and PAO1 + CV8_VAP (n = 33; intralobe, n = 18; nebulized, n = 15). Thickness values (µm) are presented as median and interquartile range (IQR) (percentiles based on weighted average definition). Grey bars include both intralobe- and nebulized-treated animals, whereas yellow bars distinguish between administration routes (intralobe: diagonal lines; nebulized: squares). Statistical analysis was performed using the Kruskal–Wallis test, and p-values are reported in Table 1. Only significant differences shown: unless indicated, comparisons *vs*. PBS + PBS; lines denote group comparisons. (**B**) Representative images from SEM (top) and CLSM (bottom) show the biofilm structure and cell viability across all treatment groups. SEM images are displayed at 1500× magnification. The red arrow indicates the biofilm, and the black arrow points to the exposed ETT surface. In the CLSM images, LIVE/DEAD BacLight™ staining was used to differentiate live (green) from dead (red) bacteria. The white arrow indicates the ETT.

**Table 1.** Statistical analysis of in *vivo*–formed ETT biofilm thickness following treatment with PBS, CV8, or CV8_VAP. Thickness values (µm) are reported as median and interquartile range (IQR), with percentiles calculated using the weighted average method. Statistical analysis was performed using the Kruskal–Wallis test, with p-values reported for the following pairwise comparisons: II) Analysis not considering the type of administration: ^a^ PBS + PBS vs. PAO1 + PBS; ^b^ PBS + PBS vs. PAO1 + CV8; ^c^ PBS + PBS vs. PAO1 + CV8_VAP; ^d^ PAO1 + PBS vs. PAO1 + CV8; ^e^ PAO1 + PBS vs. PAO1 + CV8_VAP; ^f^ PAO1 + CV8 vs. PAO1 + CV8_VAP; II) Analysis considering the type of administration: ^a^ PBS + PBS intralobe vs. PAO1 + PBS intralobe ; ^b^ PBS + PBS intralobe vs. PAO1 + CV8 intralobe; ^c^ PBS + PBS vs. PAO1 + CV8_VAP intralobe; ^d^ PAO1 intralobe + PBS vs. PAO1 + CV8 intralobe; ^e^ PAO1 + PBS intralobe vs. PAO1 + CV8_VAP intralobe; ^f^ PAO1 + CV8 intralobe vs. PAO1 + CV8 nebulized; ^g^ PAO1 + CV8 intralobe vs. PAO1 + CV8_VAP intralobe; ^h^ PAO1 + CV8_VAP intralobe vs. PAO1 + CV8_VAP nebulized; ^i^ PAO1 + CV8 nebulized vs. PAO1 + CV8_VAP nebulized. Statistically significant p-values are shown in bold.

| Analysis not considering the type of administration |  |  |  |  |  |  |  |
| --- | --- | --- | --- | --- | --- | --- | --- |
| Thickness | PBS +PBS | PAO1 + PBS | PAO1 + CV8 |  | PAO1 + CV8_VAP | P-value |  |
| MAXIMAL | 18.43 [14.51-22.58] | 10.15 [3.65-15.12] | 7.18 [4-30-15.54] | | 5.91 [4-52-8.46] | $p=0.014$<br><sup>a</sup>$p=0.020$<br><sup>b</sup>$p=0.020$<br><sup>c</sup>$p=0.002$<br><sup>d</sup>$p=0.790$<br><sup>e</sup>$p=0.067$<br><sup>f</sup>$p=0.410$ | |
| MINIMAL | 10.87 [6.82-14.44] | 5.72 [2.03-8.92] | 2.35 [1.52-4.23] | | 2.93 [1.93-4.87] | $p=0.004$<br><sup>a</sup>$p=0.044$<br><sup>b</sup>$p=0.004$<br><sup>c</sup>$p=0.004$<br><sup>d</sup>$p=0.032$<br><sup>e</sup>$p=0.078$<br><sup>f</sup>$p=0.290$ | |
| MEAN | 18.62 [11.13-18.67] | 8.18 [2.77-11.06] | 4.88 [3.05-12.22] | | 4.70 [3.48-6.85] | $p=0.009$<br><sup>a</sup>$p=0.016$<br><sup>b</sup>$p=0.005$<br><sup>c</sup>$p=0.002$<br><sup>d</sup>$p=0.184$<br><sup>e</sup>$p=0.091$<br><sup>f</sup>$p=0.890$ | |
| Analysis considering the type of administration |  |  |  |  |  |  |  |
| Thickness | PBS +PBS | PAO1 + PBS | PAO1 + CV8<br>intralobar | PAO1 + CV8<br>nebulized | PAO1 +<br>CV8_VAP<br>intralobar | PAO1 + CV8_VAP<br>nebulized | P-value |
| MAXIMAL | 18.43 [14.51-22.58] | 10.15 [3.65-15.12] | 7.38 [3.97-16.12] | 6.18 [4.49-15.06] | 7.19 [5.63-10.43] | 4.56 [3.38-5.94] | $p=0.014$<br><sup>a</sup>$p=0.020$<br><sup>b</sup>$p=0.027$<br><sup>c</sup>$p=0.004$<br><sup>d</sup>$p=0.838$<br><sup>e</sup>$p=0.293$<br><sup>f</sup>$p=0.824$<br><sup>g</sup>$p=0.874$<br><sup>h</sup>$p=0.011$<br><sup>i</sup>$p=0.042$ |
| MINIMAL | 10.87 [6.82-14.44] | 5.72 [2.03-8.92] | 1.85 [1.11-3.30] | 3.54 [2.66-7.60] | 4.26 [3.10-6.79] | 2.17 [1.53-2.50] | $p=0.001$<br><sup>a</sup>$p=0.044$<br><sup>b</sup>$p=0.005$<br><sup>c</sup>$p=0.014$<br><sup>d</sup>$p=0.011$<br><sup>e</sup>$p=0.599$<br><sup>f</sup>$p=0.020$<br><sup>g</sup>$p=0.005$<br><sup>h</sup>$p=0.001$<br><sup>i</sup>$p=0.017$ |
| MEAN | 14.62 [11.13-18.67] | 8.18 [2.77-11.06] | 4.88 [2.50-9.11] | 4.64 [3.31-10.54] | 6.08 [4.66-8.99] | 3.67 [2.82-4.45] | $p=0.004$<br><sup>a</sup>$p=0.016$<br><sup>b</sup>$p=0.004$<br><sup>c</sup>$p=0.006$ |
| | | | | | | | <sup>d</sup> $p=0.136$<br><sup>e</sup> $p=0.447$<br><sup>f</sup> $p=0.617$<br><sup>g</sup> $p=0.359$<br><sup>h</sup> $p=0.002$<br><sup>i</sup> $p=0.042$ |

With regard to the administration route, CV8 treatment produced similar effects when delivered intralobe or by nebulization. In contrast, for CV8_VAP, nebulization significantly reduced maximal, minimal, and mean biofilm thickness compared with intralobar administration. These results highlight the enhanced efficacy of the nebulized treatment with CV8_VAP in disrupting biofilm integrity, likely due to the direct contact of the nebulized CV8_VAP with the PAO1 biofilm, which may favor its persistence and activity (Figure 2A, Table 1).

We used then CLSM to qualitatively assess the structural integrity of the biofilm (Figure 2). In the PBS + PBS, PAO1 + PBS, and CV8 intralobar groups, biofilms were observed to be dense and well organized, with a robust extracellular matrix characteristic of mature biofilms and an increased proportion of viable cells (green fluorescence). In contrast, the biofilms in the CV8_VAP treated groups, particularly in the nebulized group, displayed disrupted architecture with thinner, less cohesive biofilm layers with lower bacteria viability (red fluorescence) (Figure 2B).

These results corroborate the *ex vivo* data obtained from ETTs of patients (12) and support the disruptive effect of the VAP platform (in CV2 and CV8 chassis) on *in vivo*–formed ETT biofilms, with nebulization appearing to exert a more pronounced therapeutic effect.

### Impact of CV8_VAP administration on *P. aeruginosa* viability

SEM and CLMS results suggested a decrease in bacterial load in ETTs after treatment with VAP platform (Figure 2). To support this hypothesis, we evaluated the *P. aeruginosa* PAO1 load and the concomitant cultured aerobic bacteria in ETTs and cuffs.

The presence of *P. aeruginosa* was confirmed only in the PAO1 inoculated groups (Figure 3). Animals treated with CV8_VAP (intralobe + nebulized) exhibited a non-statistically significant reduction of 0.4 log CFU/mL in *P. aeruginosa* load within the ETT compared to those receiving PBS and CV8 (Figure 3A, Table 2). A sub-analysis separating intralobe from nebulized administration indicated non-statistically significant reductions in bacterial load for the CV8_VAP nebulized group, amounting to 0.6 and 1.2 log CFU/mL relative to PBS and PAO1+CV8 nebulized controls, respectively. These findings were consistent with a trend toward lower minimal biofilm thickness in the ETT of the PAO1 + CV8_VAP nebulized group (Figure 2). By contrast, regarding *P. aeruginosa* load in the cuff, we observed a non-significant decrease in bacterial burden when treatment was administered by intralobe instillation, regardless of the treatment group. A potential explanation for this finding lies in the fact that nebulization is delivered via the lumen of ETT, and therefore only a residual effect would be expected on the cuff, which is located on the external surface of the ETT. Non-statistically significant differences in other bacterial loads in the ETT and cuffs across treatment groups are reported (Figure 3A, Table 2).

**Figure 3.**
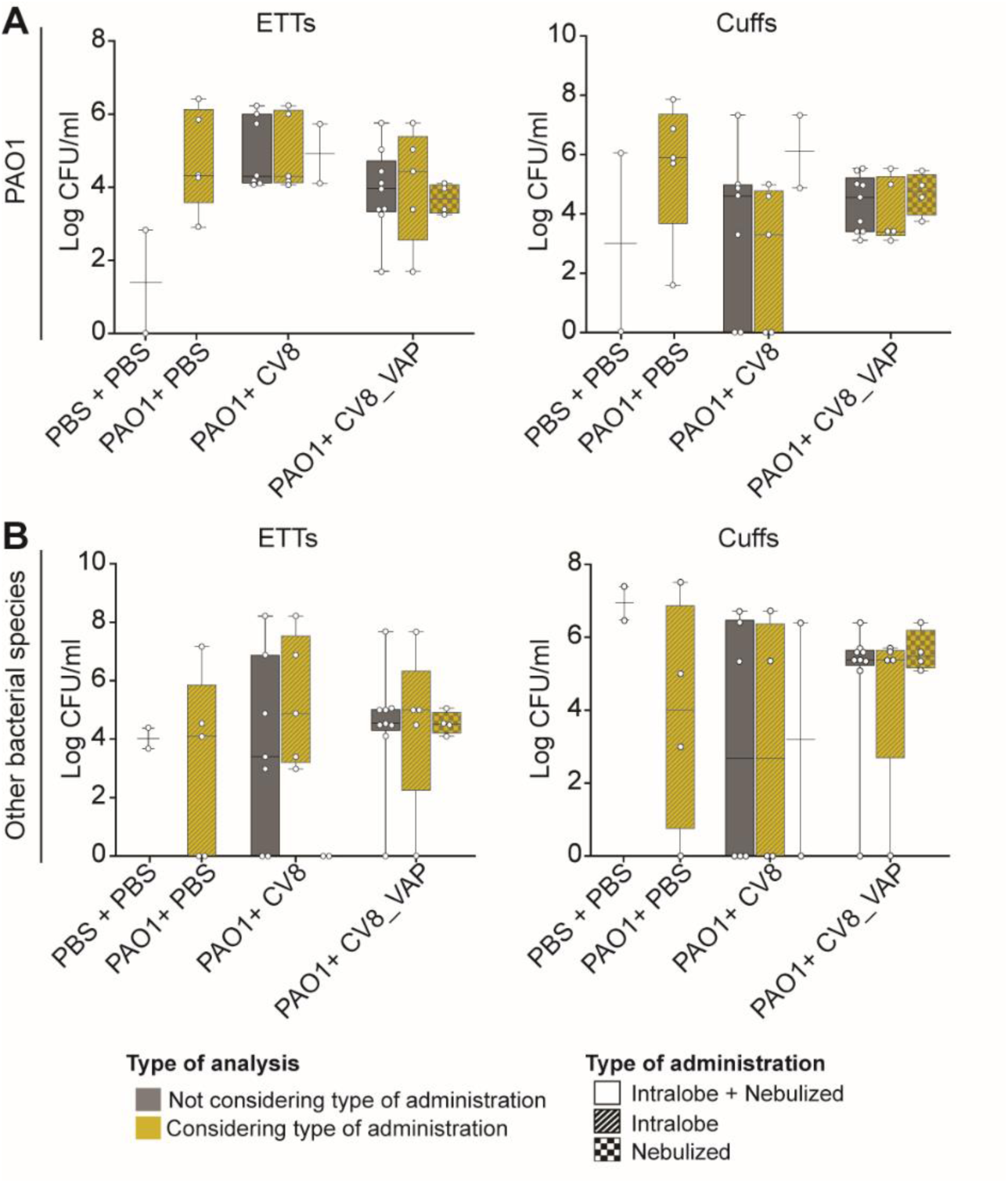
*P. aeruginosa* (A) and other bacterial species (B) burden after treatment in ETTs or cuffs. Groups were as follows: PBS + PBS (n = 2, intralobe), PAO1 + PBS (n = 5, intralobe), PAO1 + CV8 (n = 7; intralobe, n = 5; nebulized, n = 2), and PAO1 + CV8_VAP (n = 9; intralobe, n = 5; nebulized, n = 4). Grey bars include both intralobe- and nebulized-treated animals, whereas yellow bars distinguish between administration routes (intralobe: diagonal lines; nebulized: squares). Statistical analysis was performed using the Kruskal–Wallis test, and p-values are reported in Table 2. Bar plots of PBS + PBS and PAO1 + CV8 nebulized groups (n = 2) are not shown, as IQR cannot be calculated.

**Table 2.**
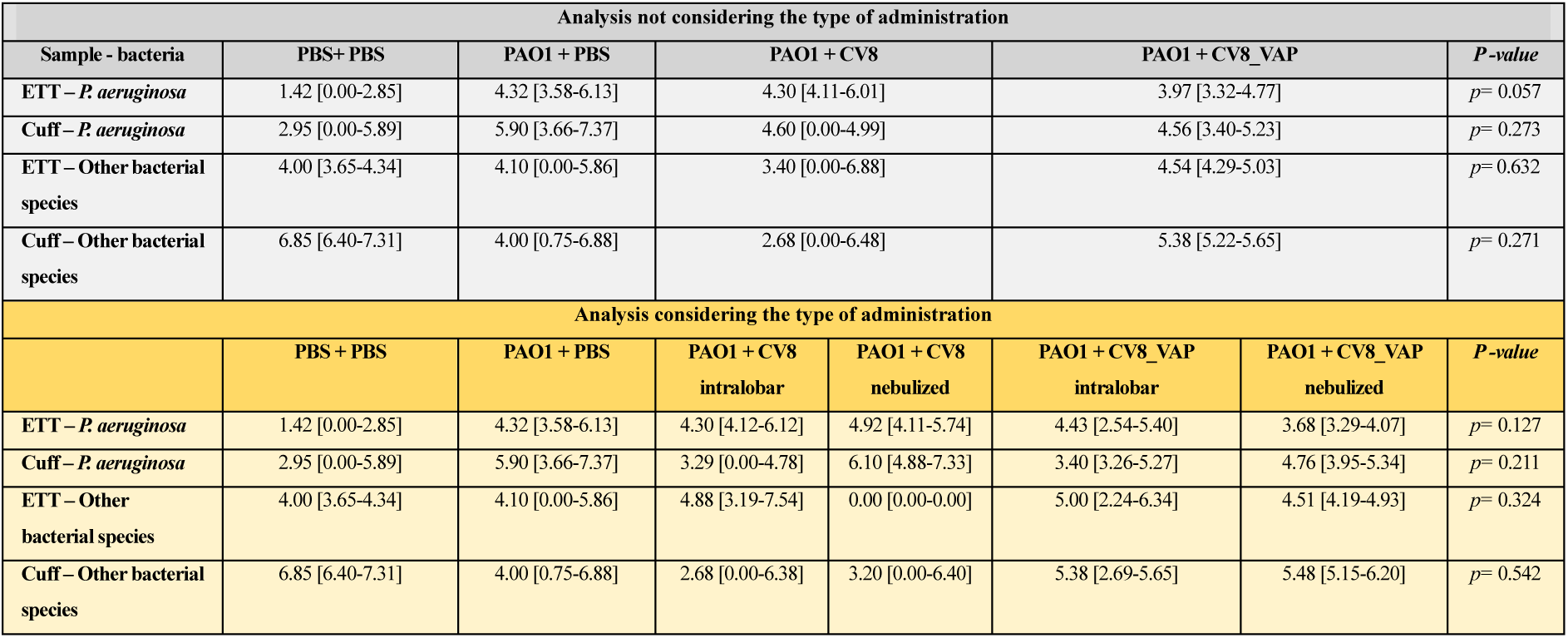
Statistical analysis of *P. aeruginosa* and other bacterial species load in ETTs and cuffs. Median and IQR values showed no statistically significant differences among groups. Percentiles were calculated using the weighted average method. Statistical analysis was performed using the Kruskal-Wallis test, with p-values indicated.

Mechanically ventilated patients rapidly lose microbial diversity after intubation, a condition further aggravated by antibiotics, increasing the risk of lower respiratory tract infections (22). Considering this, we evaluated the bacterial composition of the ETTs and cuffs after CV8_VAP platform treatment (Figure 3B, Table 2).

The biofilm in the PBS + PBS group was characterized by the presence of concomitant cultured aerobic bacteria, supporting previous reports that ETTs can be rapidly colonized by secretions, cell debris, and microorganisms originating from the stomach and oropharynx (11,23). Among the PAO1 infected groups, excluding *P. aeruginosa*, the analysis did not reveal statistically relevant differences, although we could see a slight increase in bacterial number in the cuffs group of PAO1+CV8_VAP. These data suggest that the concomitant cultured aerobic bacteria was preserved in the treated groups within ETTs or cuffs, indicating that treatment with CV8_VAP primarily affected *P. aeruginosa* viability.

Our findings suggest a trend toward a reduction in *P. aeruginosa* load and a biofilm minimal thickness within ETT biofilms in the VAP treated group, particularly when administered via nebulization.

### Impact of the VAP Platform on *P. aeruginosa-*Induced Pneumonia

Concurrently, we investigated the therapeutic potential of CV8_VAP against *P. aeruginosa*–induced pneumonia. Compared with control groups, *P. aeruginosa* load was significantly reduced in BALF and lung of the CV8_VAP-treated group, independent of administration route (Figure 4A). Although no statistically significant differences were observed between intralobe or nebulized groups, nebulization of CV8_VAP appeared to enhance the effect. CV8 and CV8_VAP showed similar lung bacterial loads, suggesting the therapeutic effect was driven by the VAP platform (Figure 4B).

**Figure 4.**
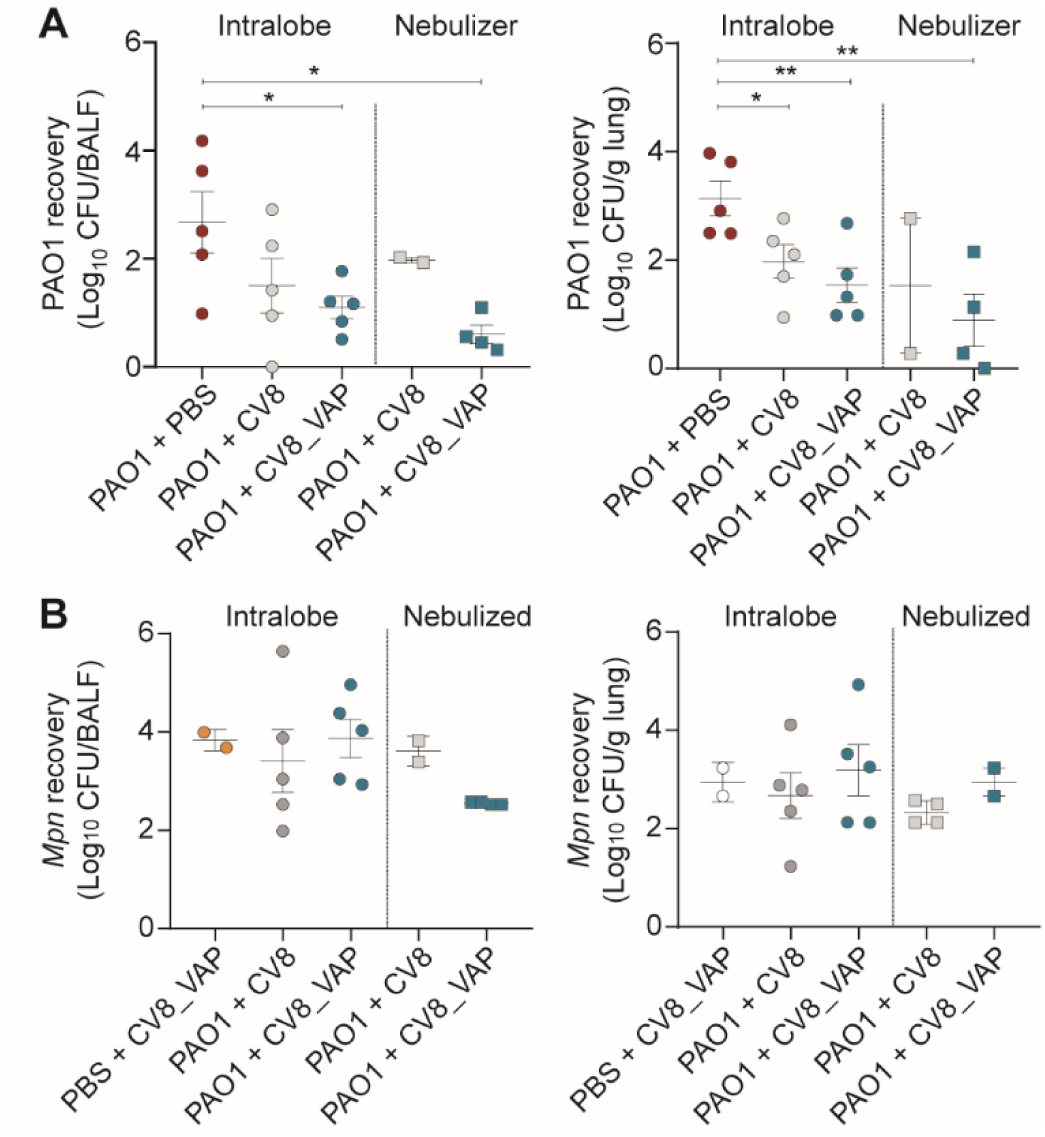
Bacterial load in the lung after treatment. Bacterial load of PAO1 (A) and *Mpn* strains (CV8 or CV8_VAP) (B) recovered from BALF (left) and lung tissue (right), shown as Mean ± SD. (B) Statistical analysis was performed using an unpaired t-test (\**p*<0.05; **p<0.01).

We then evaluated the VAP platform’s effect on *P. aeruginosa*-induced inflammation. Infection upregulated IL-1β, IL-6, IL-8, and IFN-γ expression (Figure 5), whereas CV8_VAP did not trigger any significant inflammatory response, reinforcing its safety profile observed in minipigs (Figure 1). CV8_VAP intralobe administration reduced the expression of all tested inflammatory genes to baseline levels (Figure 5A). Similar results were observed for protein levels of inflammatory markers following intralobe administration (Figure 5B). In the nebulized administration, a comparable overall effect was detected; however, in some cases either mRNA or protein levels of specific inflammatory markers were not reduced. This discrepancy suggests that the findings in nebulized animals may reflect experimental variability, or alternatively, that the administration method influences specific inflammatory pathways (e.g., IL-1β), which should be further investigated.

**Figure 5.**
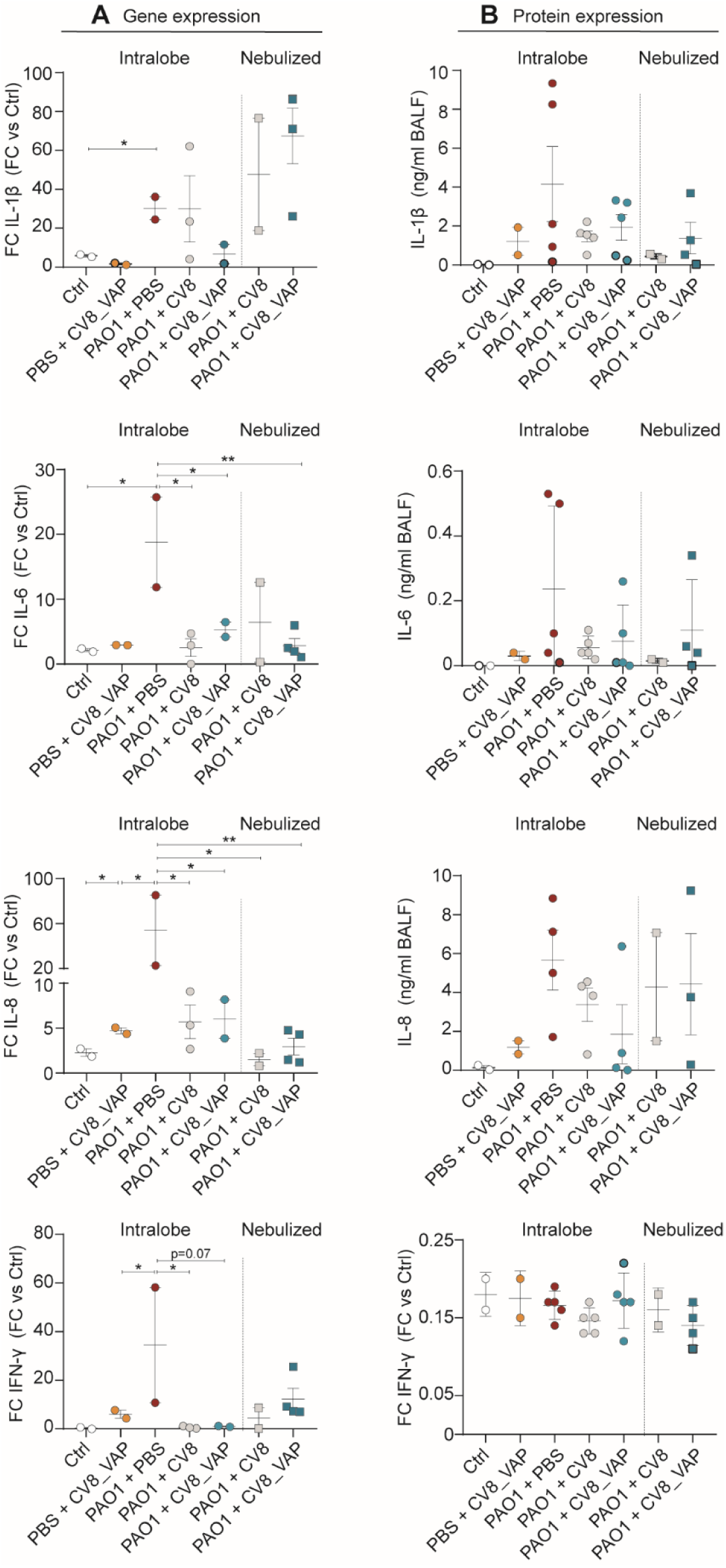
Effect of CV8_VAP in *P. aeruginosa*-induced lung inflammation. **(A)** Fold change (FC) of RNA expression of inflammatory markers in BALF samples in comparison with PBS + PBS group (FC=1). Data are shown as Mean ± SEM. Statistical analyses were performed by using unpaired t-test (*p<0.05, ** p<0.01). **(B)** Inflammatory proteins detected in BALF samples (ng/ml). Protein detection was determined by using ELISA Multiplex assay.

Surprisingly, CV8 administration reduced the expression of the inflammatory markers, suggesting that CV8 platform could have an anti-inflammatory effect, independent of the expression of the VAP platform. Notably, no systemic inflammatory response was detected (Supplementary Figure 3A-D).

Finally, lung damage was assessed based on macroscopic and histological examination (Supplementary Figure 3E-F, and Supplementary Table 3). No statistically significant differences were observed between the groups. This suggests that CV8_VAP lowers bacterial load and inflammation, but does not improve lung tissue damage, possibly due to prior *P. aeruginosa* infection or intrinsic injury associated with mechanical ventilation.

### Microbiome composition in ETT-biofilms and lung

As previously noted, CV8_VAP showed a specific effect against PAO1. However, potential shifts in the microbial community composition could not be excluded. To explore this, we analyzed the taxonomic diversity of biofilm-associated microbial communities in ETTs and lung tissue (Figure 6).

**Figure 6.**
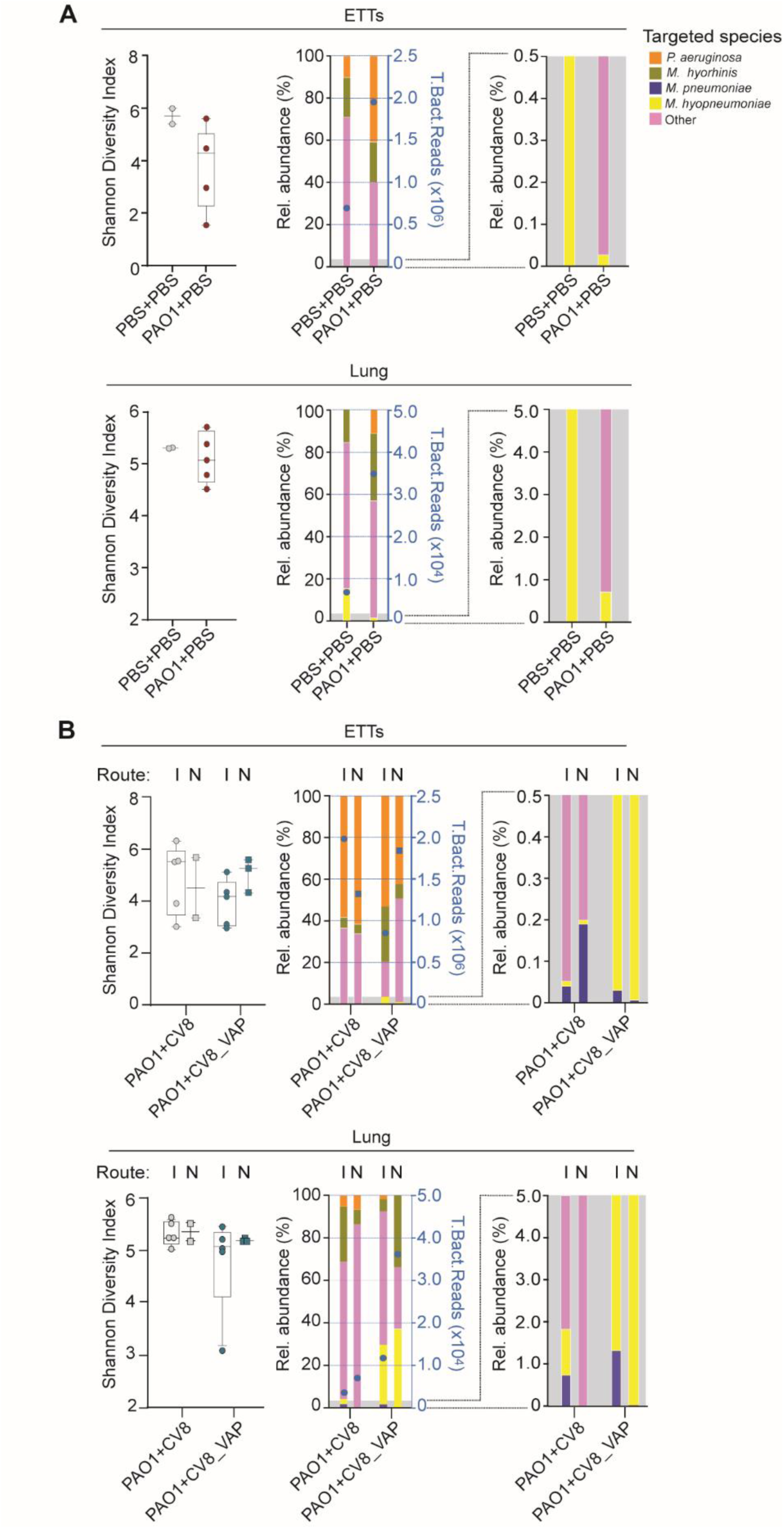
Impact of the VAP platform on ETT and lung microbiome after PAO1 infection in control (A) and CV8 or CV8_VAP treated groups (B). Left, Shannon diversity index. Data is presented as median and IQR, with percentiles calculated using the weighted average method. Bar plots of PBS + PBS and PAO1 + CV8 nebulized groups (n = 2) are not shown, as IQR cannot be calculated. Statistical analysis was performed using the Kruskal-Wallis test, with results shown in Table 3. **Right,** Relative abundance of top taxa identified in ETTs both in intralobe (I) and nebulized (N) animals. Left Y-axes (black) show relative abundance; right Y-axis (blue) represents total bacterial reads for the selected species, including *P. aeruginosa*, *M. hyorhinis*, *M. hyopneumoniae*, *M. pneumoniae*, and others (all remaining taxa with relative abundances lower than those of the top five taxa). A close-up of species with a relative abundance of less than 0.5 % is shown in the lower right corner. Taxa names are shown at the highest taxonomic resolution available, including genus and species when possible. Animals included: PBS + PBS (n = 2; intralobe), PAO1 + PBS (n = 5; intralobe), PAO1 + CV8 (n = 7; intralobe, n = 5; nebulized, n = 2), and PAO1 + CV8_VAP (n = 9; intralobe, n = 5; nebulized, n = 3). Pig 544 is not reported because of the low quality of the DNA sample that compromised the results.

First, we compared the control groups PBS + PBS and PAO1 + PBS. In ETTs, a decrease in the Shannon diversity index (SDI) indicated reduced microbiome diversity when comparing biofilms from healthy pigs (PBS + PBS; n = 2; SDI 5.70 [5.40–6.01]) with those challenged with *P. aeruginosa* (n = 4; SDI 4.32 [2.29–5.06]) (Figure 6A, upper panel; Supplementary Figure 4; Table 3). Relative abundance analysis based on 16S rRNA sequencing revealed that bacteria representing <0.5% relative abundance (classified as ‘other bacteria’) were replaced by PAO1. Furthermore, a significant anti-correlation was observed between *P. aeruginosa* and *Mycoplasma hyopneumoniae* (r = –0.49, p = 0.02), a commensal bacterium of the porcine respiratory tract (Figure 6A). In the lung, this shift was associated with a modest decrease in SDI in the PAO1 + PBS group (5.06 [4.65–5.64]) compared with the PBS group (5.30 [5.29–5.31]) (Figure 6A, bottom panel; Table 3). This effect was less pronounced than in the ETTs, likely reflecting limited migration of *P. aeruginosa* from the ETT to the lower airways, consistent with its relative abundance in lung tissue (Figure 6B, Supplementary Figure 4). As in the ETTs, the presence of PAO1 was linked to a decrease in *M. hyopneumoniae*, suggesting that colonization by this pathogen drives a shift in the natural microbiota of both upper and lower airways.

**Table 3.** Statistical analysis Shannon Diversity Index of top taxa identified in ETT and lung after microbiome sequencing. Shannon diversity index values are shown as median and interquartile range (IQR), with percentiles calculated using the weighted average method. Statistical analysis was performed using the Kruskal-Wallis test, with p-values indicated.

| ETTs |  |  |  |  |  |  |
| --- | --- | --- | --- | --- | --- | --- |
| Analysis not considering the type of administration |  |  |  |  |  |  |
| PBS+PBS | PAO1 + PBS | PAO1 + CV8 |  | PAO1 + CV8_VAP |  | p-value |
| 5.70 [5.40-6.01] | 4.32 [2.29-5.06] | 4.11 [2.49-4.22] |  | 3.50 [2.51-4.31] |  | p= 0.161 |
| Analysis considering the type of administration |  |  |  |  |  |  |
| PBS+PBS | PAO1 + PBS | PAO1 + CV8 intralobar | PAO1 + CV8 nebulized | PAO1+CV8_VAP intralobar | PAO1 + CV8_VAP nebulized | p-value |
| 5.70 [5.40-6.01] | 4.32 [2.29-5.06] | 4.11 [2.57-4.41] | 3.36 [2.49-4.22] | 3.11 [2.25-3.52] | 4.48 [4.12-4.60] | p= 0.147 |
| Lung |  |  |  |  |  |  |
| Analysis not considering the type of administration |  |  |  |  |  |  |
| PBS+PBS | PAO1 + PBS | PAO1 + CV8 |  | PAO1 + CV8_VAP |  | p-value |
| 5.30 [5.29-5.31] | 5.06 [4.65-5.64] | 5.23 [5.15-5.51] |  | 5.17 [5.02-5.22] |  | p= 0.356 |
| Analysis considering the type of administration |  |  |  |  |  |  |
| PBS+PBS | PAO1 + PBS | PAO1 + CV8 intralobar | PAO1 + CV8 nebulized | PAO1+CV8_VAP intralobar | PAO1 + CV8_VAP nebulized | p-value |
| 5.30 [5.29-5.31] | 5.06 [4.65-5.64] | 5.23 [5.12-5.55] | 5.32 [5.15-5.49] | 5.04 [4.05-5.33] | 5.18 [5.16-5.22] | p= 0.595 |

We next examined the ETT microbiome composition in the PAO1 infected animals treated with CV8- and CV8_VAP. In both cases, the SDI was reduced, as observed in the PAO1 + PBS group, irrespective of administration route (Figure 6B, upper panel; Table 3). This reduction was associated with a decrease in bacteria representing <0.5% relative abundance (‘other bacteria’). The presence of *M. pneumoniae* was also confirmed in these groups (Figure 6B, upper panel; Supplementary Figure 4; Table 3), although no significant correlations were detected between *M. pneumoniae* and *P. aeruginosa*. Interestingly, the CV8-treated group displayed a bacterial composition similar to the infected control (PAO1 + PBS), whereas in CV8_VAP-treated animals, an increase in *M. hyopneumoniae* was observed under both intralobar and nebulization routes. This profile, more comparable to the PBS + PBS group, suggests recolonization of the ETT by commensal bacteria. Supporting this hypothesis, a similar trend was observed in lung tissue (Figure 6B, bottom panel; Table 3).

In conclusion, CV8_VAP treatment not only reduced PAO1 pathogenic biofilms and lung bacterial load, but also helped preserve or restore microbial diversity in both ETT biofilms and pulmonary tissue, as reflected by increased abundance of *M. hyopneumoniae* and *M. hyorhinis*.

### Clinical outcome of VAP treated animals

Ventilator parameters showed similar dynamics across all groups (Figure 7). Animals required progressively higher FiO₂ (Figure 7A), PEEP (Figure 7B), and RR (Figure 7C) due to worsening oxygenation, corroborated by a significant decline in PaO₂/FiO₂ ratio over time (Figure 7D). Respiratory mechanics also deteriorated, with significant increases in driving (Figure 7E) and peak airway pressures (Figure 7F), and a decline in lung compliance (Figure 7G). Notably, peak airway pressure was significantly higher in PAO1+CV8_VAP *vs*. PBS+CV8_VAP (*p*=0.02), and lung compliance was reduced in both PAO1+PBS (*p*=0.04) and PAO1+CV8_VAP (*p*=0.02) compared to PBS. Among nebulized groups, driving pressure was significantly worse in CV8-treated animals.

**Figure 7.**
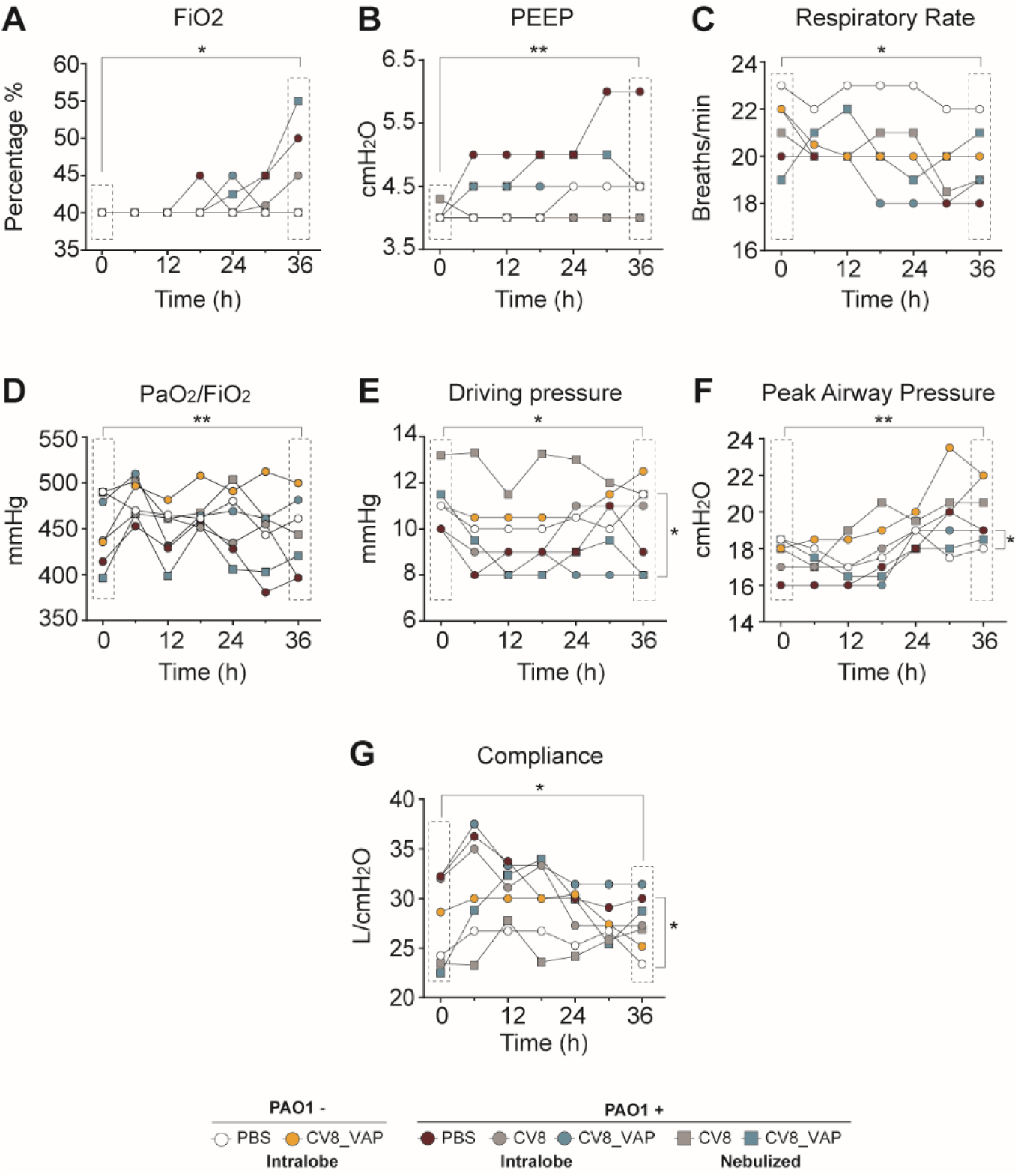
Time-course evaluation of ventilatory parameters across treatment groups. Animals were grouped according to treatment: PBS (open symbols), PBS + CV8_VAP (orange), PAO1 + PBS (red), PAO1 + CV8 (grey), and PAO1 + CV8_VAP (blue). Administration routes are indicated by symbol shape: intralobe (circles) and nebulized (squares). Data are presented as median values over time for the following parameters: **(A)** Fraction of Inspired Oxygen (FiO₂, %), **(B)** Positive End-Expiratory Pressure (PEEP, cmH_2_O), **(C)** Respiratory Rate (breaths/min), **(D)** Oxygenation Index (PaO₂/FiO₂ ratio, mmHg), **(E)** Driving Pressure (mmHg), **(F)** Peak Airway Pressure (cmH_2_O), **(G)** Lung Compliance (L/cmH_2_O). Statistical analysis was performed using two-way ANOVA followed by post-hoc multiple comparisons with false discovery rate (FDR) correction (*p< 0.05; **p< 0.001).Significant differences over time are indicated by dashed boxes, while group-wise comparisons at specific time points are shown on the right side of each panel.

Hemodynamic variables varied significantly over time. MAP initially declined at 6 h and partially recovered by study end (Figure 8A), while PCWP increased steadily, suggesting early cardiac dysfunction (Figure 8B). Between-group comparisons revealed significant differences in MAP, PCWP, and mPAP (Figure 8A–C). Post-hoc analysis showed significantly lower mPAP and PCWP in the PBS group *vs*. PAO1-infected groups (Table 5), indicating increased pulmonary vascular resistance and cardiac preload. MAP was significantly reduced in PAO1+CV8_VAP animals versus PBS+PBS (p=0.049) and PBS+CV8_VAP (p=0.035), pointing to a potential hypotensive effect requiring further investigation. No significant differences were found among treatment groups.

**Figure 8.**
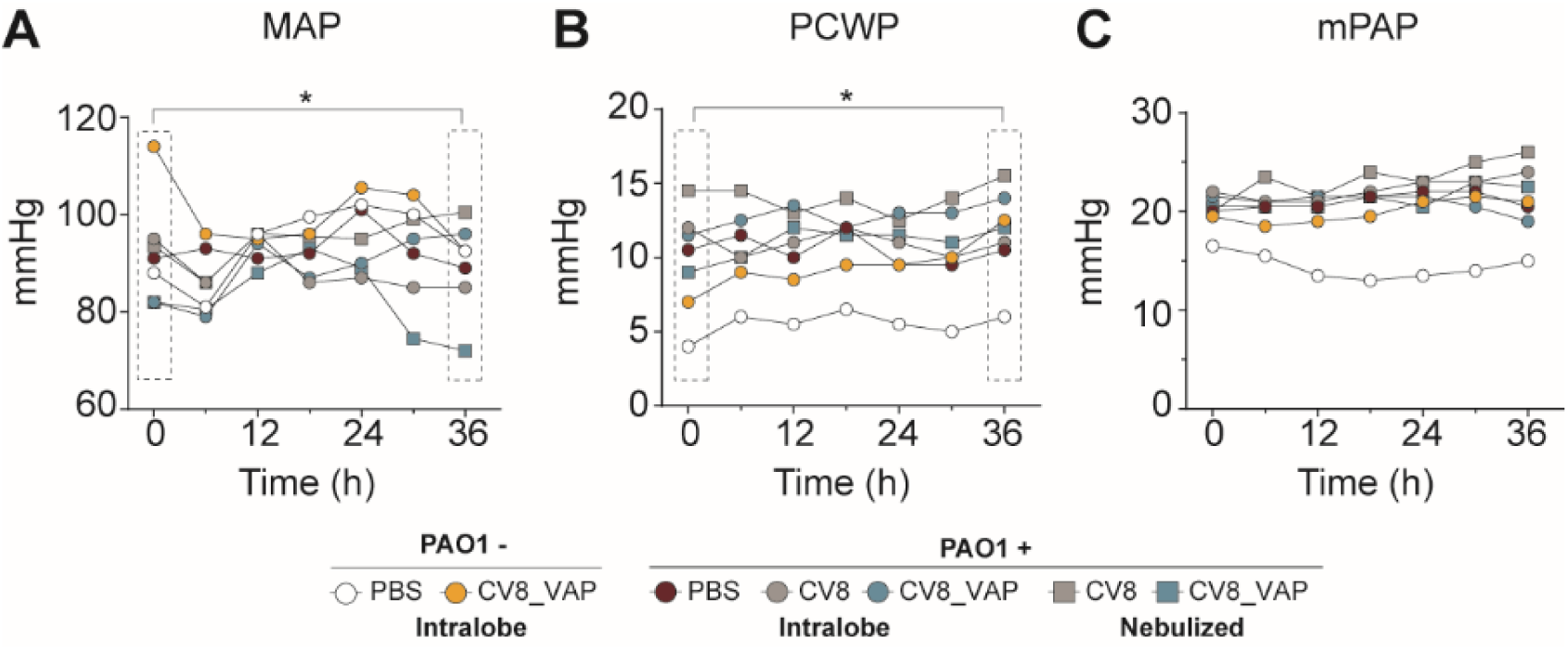
Hemodynamic parameters over time across experimental groups. Animals were grouped according to treatment: PBS (open), PBS + CV8_VAP (orange), PAO1 + PBS (red), PAO1 + CV8 (grey), PAO1 + CV8_VAP (blue). Intralobe (circles) and nebulized (squares) administration routes are shown. Data are presented as median values for the following parameters: **(A)** Mean Arterial Pressure (MAP, mmHg), **(B)** Pulmonary Capillary Wedge Pressure (PCWP, mmHg), and **(C)** Mean Pulmonary Arterial Pressure (mPAP, mmHg). Statistical analysis was performed using two-way ANOVA followed by post hoc multiple comparisons with false discovery rate (FDR) correction (*p< 0.05; **p< 0.001). Significant differences over time are indicated by dashed boxes.

Biochemical and hematological analyses (Supplementary Table 4) revealed significant between-group differences in WBC counts, though not over time. Post-hoc results showed WBC counts were lower in PBS+CV8_VAP, PAO1+CV8, and PAO1+CV8_VAP compared to PBS, suggesting possible modulation of the immune response. Among infected groups, PAO1+PBS animals had significantly higher WBC counts than those receiving CV8 or CV8_VAP, supporting a treatment-associated reduction in systemic inflammation.

Creatinine and platelet counts showed significant temporal variation, with mild creatinine increases and declining platelets, consistent with early renal dysfunction and thrombocytopenia typical of systemic inflammation or sepsis. Liver enzymes (ALT, GGT, ALP) fluctuated, with ALP showing both group- and time-dependent differences; post-hoc analysis indicated lower ALP levels in animals treated via nebulization. Coagulation parameters also changed over time, without intergroup differences (Supplementary Table 4).

Together, these findings demonstrate that infection with *P. aeruginosa* induces marked systemic and pulmonary deterioration, affecting respiratory function, hemodynamics, and multiple organ systems. Importantly, the CV8_VAP platform, particularly when administered via nebulization, showed signs of mitigating the inflammatory and organ dysfunction responses associated with CV8_VAP, supporting its therapeutic potential.

### Deep into the nebulized treatment

Both intralobe and nebulized delivery of CV8_VAP showed therapeutic benefit, but nebulized administration is more clinically relevant and translatable. To evaluate safety, we compared key clinical outcomes between animals treated with nebulized CV8 and CV8_VAP.

Nebulization lasted 55–75 minutes in all animals. While not statistically significant, trends indicated that CV8-treated animals had higher peak airway pressures (Figure 9A) and sustained elevations in driving pressure (Figure 9B), suggesting increased respiratory resistance and reduced lung compliance. MAP values were comparable (Figure 9C), though slightly more stable in the CV8_VAP group. Heart rate was also similar (Figure 9D), with greater variability observed in CV8-treated animals, possibly reflecting higher physiological stress.

**Figure 9.**
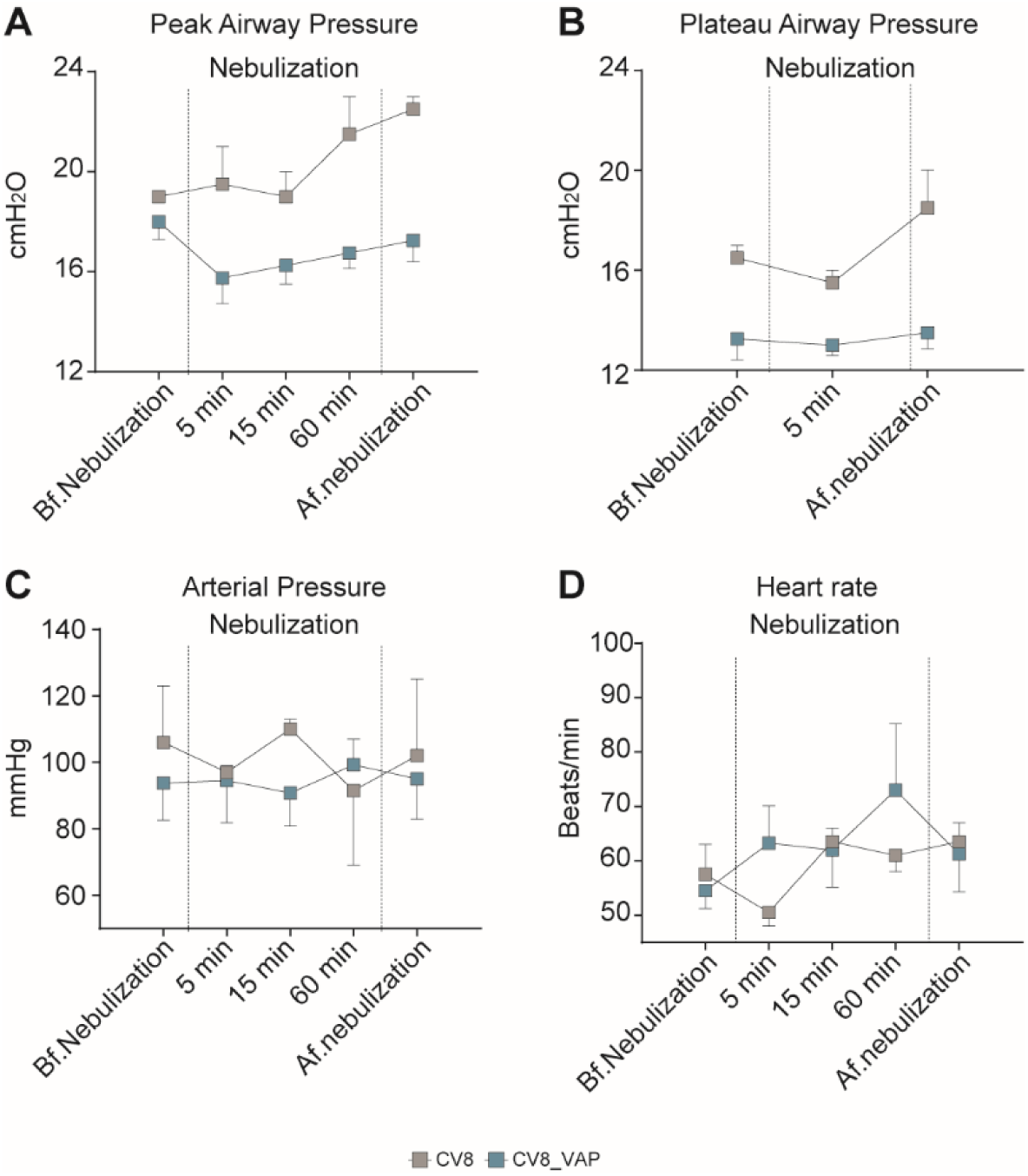
Clinical outcomes in animals treated with nebulized CV8 (grey) or CV8_VAP (blue). Median values for each group are shown across time points for the following parameters, recorded during before (Bf), the nebulization protocol and after (Af): **(A)** Peak Airway Pressure (cmH_2_O), **(B)** Plateau Airway Pressure (cmH_2_O), **(C)** Mean Arterial Pressure (mmHg), and **(D)** Heart Rate (beats/min).

Despite more stable ventilator and hemodynamic parameters with CV8_VAP, isolated adverse events occurred. One PAO1+CV8_VAP animal experienced hemodynamic decompensation requiring atropine, and another excluded animal died shortly after PAO1+CV8_VAP nebulization due to anaphylaxis, despite resuscitation efforts (Supplementary Table 5).

In conclusion, CV8_VAP improved respiratory mechanics and hemodynamic stability, supporting its therapeutic potential in *P. aeruginosa* lung infection. However, serious adverse events underscore the need for protocol refinement and close monitoring in future translational studies.

## Discussion

VAP is the most severe infectious complication in intubated and mechanically ventilated patients. It increases the duration of mechanical ventilation, ICU and hospital stays, and crude mortality.

The pathophysiology of VAP is multifactorial, involving mechanical disruption of bronchial defenses, aspiration of oropharyngeal and gastric contents, and impaired immune responses. One of the earliest events in the development of VAP is the formation of biofilms within ETTs, often within the first 24 hours after intubation. These biofilms serve as reservoirs for microorganisms, shielding them from antibiotic penetration and thereby reducing the efficacy of antimicrobial treatment. Furthermore, fragments of the biofilm can be dislodged and propelled into the lower respiratory tract by the positive pressure generated during mechanical ventilation, leading to colonization of the lower airways and potential compromise of pulmonary function.

Broad-spectrum antibiotics, while used to manage infections, contribute to microbiome dysbiosis, loss of commensals, and overgrowth of resistant pathogens such as *P.aeruginosa.* MDR strains are increasingly prevalent and pose significant therapeutic challenges. Reducing the use, the number (one vs two or more) and duration of antibiotic treatments is essential to mitigate the global MDR/XDR crisis (24). In addition, a recent study published by our group suggested that the empirical use of two antipseudomonal antibiotics in VAP patients without shock was followed by a trend to higher mortality (9).

In this context, we investigated a novel non-antibiotic therapy using a live bioengineered *Mycoplasma pneumoniae* strain expressing anti-biofilm and bactericidal agents (PelAh, PslGh, A1-II’ alginate lyase, and pyocin L1) (CV8_VAP platform). We evaluated the effect of CV8_VAP on *P. aeruginosa* PAO1 biofilms formed *in vivo*, as well as its impact on the ETT biofilm microbiome when administered as a treatment in a porcine model of VAP induced by ceftriaxone-resistant *P. aeruginosa*.

CV8_VAP significantly reduced *P. aeruginosa* burden on ETT surfaces, with no similar effects on other bacterial species, indicating pathogen specificity. Biofilm thickness and viability, assessed via SEM and CLSM, confirmed reduced biofilm formation in the CV8_VAP group.

One of the major findings was that CV8_VAP, when delivered by bronchoscopy instillation, significantly reduced *P. aeruginosa* load in both BALF and lung tissue compared to CV8 and PBS controls, confirming previous results in murine models (12). Interestingly, the lung microbiome was not markedly altered after 12 h of CV8_VAP administration, while a restoration of commensal bacteria (such as *M. hyopneumoniae*) was observed in both ETTs and lung tissue. This is an important point since the preservation of lung microbiome is directly associated with better outcomes in patients mechanically ventilated and VAP. Further reductions in bacterial burden were observed when CV8_VAP was delivered via nebulization instead of direct instillation, suggesting improved lung distribution. *P. aeruginosa* lung clearance was accompanied by reduced levels of inflammatory cytokines, indicating an attenuation of local lung inflammation, and showed consistent trends across both delivery methods although with some discrepancies between mRNA and protein levels in the nebulized animals. Despite reductions in bacterial burden and downregulation of cytokines, total lung histopathology scores did not differ between groups. This may indicate that the 12-hour treatment was too short to detect histological changes, especially considering prior murine studies where improvements were evident after 14 days post-infection. Interestingly, the fact that we see also a significant effect of CV8 in many of the parameters measured in the lung could be due to host response to antigen, or the innate DNAse and protease activity of *M. pneumoniae* (25), which could help dissolve biofilms, as well as the possibility that some of the small ORFs identified as potential antimicrobial peptides could affect *P. aeruginosa* (26).

VAP is typically treated with intravenous (IV) antibiotics that reach the alveoli via the systemic circulation. However, bacterial colonization can persist in the bronchi despite IV antibiotic therapy, contributing to relapses and recurrent episodes of VAP. While nebulized antibiotics may not effectively penetrate the consolidated regions of the lung, they achieve high concentrations along the inner surface of the ETT and in both the upper and lower bronchi. This distribution pattern likely contributed to the improved outcomes observed in our study with nebulized administration. Nevertheless, the combination of CV8_VAP with IV antibiotics will be explored further to assess potential synergistic effects.

Despite promising results, several limitations must be acknowledged. The animals used were healthy and lacked comorbidities common in VAP patients, limiting generalizability. The relatively short experimental duration (36– 72 hours) may not have captured long-term effects on lung histology or microbiome dynamics. Sample sizes, though consistent with other large-animal studies, may limit detection of subtle differences. Lastly, although intralobe instillation was employed, this method is not commonly used clinically—unlike nebulization, which offers a more translatable delivery route.

Various strategies have been explored to reduce or eliminate biofilms in preclinical and clinical settings, including systemic antimicrobials with good airway penetration (27,28), antibiotic- or silver-coated ETTs , and nebulized antibiotics to achieve high local concentrations. Mechanical approaches like the mucus shaver have also been investigated (29). While many of these methods show efficacy, their clinical use is often limited by practical challenges such as the need for extubation and reintubation, risk of VAP recurrence, and high costs. In this context, our strategy offers distinct advantages. CV8_VAP effectively reduces *P. aeruginosa* biofilm formation and acts through a non-antibiotic mechanism, potentially minimizing resistance development. Its modular, pathogen-adaptable design and compatibility with nebulization make it a practical and versatile option for clinical use. CV8_VAP could be used alongside antibiotics to treat *P. aeruginosa* VAP, potentially reducing antibiotic duration and resistance pressure, or sequentially after antibiotics to prevent relapse—common in clinical practice.

Our findings support that CV8_VAP is an innovative alternative that warrants further clinical investigation, with large-animal safety and efficacy data paving the way for human Phase I trials.

## Acknowledgment

The project that gave rise to these results has received funding from “la Caixa” Foundation under the grant agreement HR18-00058, and the European Research Council under the European Union’s Horizon 2020 research and innovation program (670216). We thank the Spanish Ministry of Economy, Industry and Competitiveness for supporting the EMBL partnership, the Centro de Excelencia Severo Ochoa, and the CERCA Program from the Generalitat de Catalunya. Additional support was provided CB 06/06/0028/CIBER de Enfermedades Respiratorias (CIBERES), ICREA Academia / Institució Catalana de Recerca i Estudis Avançats, IDIBAPS (2.603), and the SGR 01148 (2021) program of the Generalitat de Catalunya to LFB. A.M. has received funding from the European Respiratory Society and the European Union’s H2020 research and innovation programme under the Marie Sklodowska-Curie (847462).

We also thank Enric Barbeta, Jordi Vila, Gianluigi Li Bassi, Laura Muñoz, Carla Speziale, Edoardo Forin, Sofia Misuraca, Federica Piedepalumbo, Minlan Yang, Hua Yang, Amadeo Guzzardella, and Francesco Bindo for their technical support.

Artificial intelligence (AI)–assisted technologies, specifically OpenAI’s ChatGPT, were employed during the preparation of this manuscript. These tools were used exclusively to enhance the clarity, readability, and grammatical accuracy of the text. All substantive content, analysis, and interpretation reflect the authors’ original work and expertise.

## Author Contributions

LS and AT were responsible for funding acquisition. IRA led the manuscript writing, final figure generation, and coordination of the study. LFB, AM, LS, IRA, and AT contributed to conceptualization, methodology, investigation, visualization, project administration, supervision, and writing of the original draft. IRA, LFB and GM performed the statistical analyses related to biofilm and microbiome data. IRA, CS, AS and LFB contributed to the microbiological procedures as well as the conceptualization and editing of the figures. CS, RM, KK, RC, FG, BL, NV, DC, and MA contributed to the investigation by performing *in vitro* experiments and reviewed and edited the manuscript. MR, AB contributed to methodology and critical revision of the manuscript.

