## Supplementary material for "Targeting *Pseudomonas aeruginosa* Ventilator-associated pneumonia with a non-antibiotic and biolfilm-disrupting Live Biotherapeutic: Preclinical Safety and Efficacy study"

**Table of contents**

**SUPPLEMENTARY TABLES**………………………………………………………………………..…1

Supplementary Table 1. Summary of pigs used in the Safety and Efficacy studies

Supplementary Table 2. Histopathological analysis of lung tissue following PBS or CV8 administration.

Supplementary Table 3. Histopathological lesions detected in lung tissue after treatment.

Supplementary Table 4. Hematological and biochemical parameters over time.

Supplementary Table 5. Ventilator setting and adverse events during nebulization.

**SUPPLEMENTARY FIGURES**…………………………………………………………………..…..….7

Supplementary Figure 1. Slicing and sampling of the ETT and cuffs.

Supplementary Figure 2. *In vitro* characterization of the CV8_VAP strain.

Supplementary Figure 3. Impact of the VAP Platform on P. aeruginosa-Induced lung inflammation and pulmonary injury.

### SUPPLEMENTARY TABLES

**Supplementary Table 1. Summary of pigs used in the Safety and Efficacy sub-studies.** Details of the pig strain and treatment administration route are provided.

| Purpose/Pig | Pig ID | Strain | Treatment administration |
|---|---|---|---|
| Safety/ Miniature pig | 482 | PBS (left)/CV8 (right) | Intralobe |
| | 483 | PBS (left)/CV8 (right) | Intralobe |
| | 487 | PBS | Intralobe |
| | 488 | PBS | Intralobe |
| Efficacy/Large White-Landrace x Duroc pigs | 530 | PBS | Intralobe |
| | 531 | PBS | Intralobe |
| | 497 | PAO1 + PBS | Intralobe |
| | 498 | PAO1 + PBS | Intralobe |
| | 510 | PAO1 + PBS | Intralobe |
| | 512 | PAO1 + PBS | Intralobe |
| | 513 | PAO1 + PBS | Intralobe |
| | 499 | PAO1 + CV8_VAP | Intralobe |
| | 500 | PAO1 + CV8_VAP | Intralobe |
| | 501 | PAO1 + CV8_VAP | Intralobe |
| | 514 | PAO1 + CV8_VAP | Intralobe |
| | 515 | PAO1 + CV8_VAP | Intralobe |
| | 535 | PAO1 + CV8_VAP | Nebulized |
| | 537 | PAO1 + CV8_VAP | Nebulized |
| | 543 | PAO1 + CV8_VAP | Nebulized |
| | 544 | PAO1 + CV8_VAP | Nebulized |
| | 503 | PAO1 + CV8 | Intralobe |
| | 507 | PAO1 + CV8 | Intralobe |
| | 508 | PAO1 + CV8 | Intralobe |
| | 521 | PAO1 + CV8 | Intralobe |
| | 525 | PAO1 + CV8 | Intralobe |
| | 547 | PAO1 + CV8 | Nebulized |
| | 551 | PAO1 + CV8 | Nebulized |
| | 522 | PBS + CV8_VAP | Intralobe |
| | 523 | PBS + CV8_VAP | Intralobe |

**Supplementary Table 2. Histopathological analysis of lung tissue following PBS or CV8 administration.** Score of RCr, RM, RC, LCr, LC lobes are shown. Pigs 482 and 483 were administered with PBS (left lung) and CV8 (right lung).

| | | Lung Lobe | | | | | | |
|---|---|---|---|---|---|---|---|---|
| **Group** | **Pig ID** | **RCr** | **RM** | **RCr** | **LCr** | **LC** | **Average/lung** | **PBS vs CV8 p-value** |
| PBS | 482 | N/A | N/A | N/A | 0,71 | 0,71 | **0,71** | 0.41 |
| | 483 | N/A | N/A | N/A | 0,64 | 0,43 | **0,54** | |
| | **Average/lobe** | **N/A** | **N/A** | **0,68** | **N/A** | **0,57** | **0,63** | |
| CV8 | 482 | 0,43 | 0,43 | 0,57 | N/A | N/A | **0,48** | |
| | 483 | 1,29 | 0,5 | 0,71 | N/A | N/A | **0,83** | |
| | 487 | 0,43 | 0,14 | 0,14 | 0,14 | 0,14 | **0,20** | |
| | 488 | 0,21 | 0,14 | 0,14 | 0,29 | 0,43 | **0,24** | |
| | **Average/lobe** | **0,59** | **0,30** | **0,39** | **0,21** | **0,29** | **0,44** | |

**Supplementary Table 3. Histopathological lesions detected in lung tissue after treatment.** Lesions were quantified in the right (RCr, RM, RL) and left (LCr, LC) lobes. Final score was calculated as the average across all lung lobes. Score: 0 = no pneumonia; 1 =purulent mucous plugging; 2= bronchiolitis; 3= pneumonia; 4= confluent pneumonia; 5= abscessed pneumonia.

| Administration route | Group | Pig ID | Lobe | | | | | Average/lung |
|---|---|---|---|---|---|---|---|---|
| | | | RCr | RM | RCr | LCr | LC | |
| Intralobe | PBS + PBS | 526 | 3 | 4 | 3 | 3 | 4 | **3,4** |
| | | 530 | 2 | 3 | 3 | 2 | 4 | **2,8** |
| | | 531 | 1 | 1 | 3 | 3 | 4 | **2,4** |
| | | **Average/lobe** | **3,0** | **4,0** | **3,0** | **3,0** | **4,0** | **2,9** |
| | PBS + CV8_VAP | 522 | 2 | 3 | 4 | 3 | 4 | **3,2** |
| | | 523 | 2 | 3 | 3 | 3 | 3 | **2,8** |
| | | **Average/lobe** | **2,0** | **3,0** | **3,5** | **3,0** | **3,5** | **3,0** |
| | PAO1 + PBS | 496 | 1 | 3 | 3 | 3 | 3 | **2,6** |
| | | 497 | 2 | 3 | 4 | 3 | 3 | **3,0** |
| | | 498 | 1 | 3 | 4 | 3 | 3 | **2,8** |
| | | 510 | 3 | 4 | 4 | 3 | 5 | **3,8** |
| | | 512 | 3 | 4 | 3 | 3 | 3 | **3,2** |
| | | 513 | 3 | 4 | 3 | 0 | 3 | **2,6** |
| | | **Average/lobe** | **2,2** | **3,5** | **3,5** | **2,5** | **3,3** | **3,0** |
| | PAO1 + CV8 | 503 | 2 | 2 | 3 | 2 | 4 | **2,6** |
| | | 507 | 2 | 2 | 3 | 2 | 4 | **2,6** |
| | | 508 | 0 | 3 | 3 | 3 | 4 | **2,6** |
| | | 521 | 3 | 4 | 3 | 1 | 1 | **2,4** |
| | | **Average/lobe** | **1,8** | **2,8** | **3,0** | **2,0** | **3,3** | **2,6** |
| | PAO1 + CV8_VAP | 499 | 2 | 4 | 2 | 3 | 3 | **2,8** |
| | | 500 | 2 | 3 | 3 | 1 | 3 | **2,4** |
| | | 501 | 3 | 4 | 4 | 3 | 4 | **3,6** |
| | | 514 | 1 | 3 | 2 | 4 | 3 | **2,6** |
| | | 515 | 2 | 4 | 3 | 3 | 3 | **3,0** |
| | | 525 | 2 | 3 | 4 | 3 | 3 | **3,0** |
| | | **Average/lobe** | **2,0** | **3,5** | **3,0** | **2,8** | **3,2** | **2,9** |
| Nebulized | PAO1 + CV8 | 547 | 2 | 4 | 3 | 3 | 3 | **2,8** |
| | | 550 | 2 | 4 | 3 | 3 | 5 | **3,1** |
| | | **Average/lobe** | **2,0** | **3,8** | **2,8** | **2,8** | **3,5** | **3,0** |
| | PAO1 + CV8_VAP | 535 | 4 | 4 | 4 | 4 | 4 | **4,0** |
| | | 537 | 4 | 4 | 4 | 4 | 3 | **3,5** |
| | | 543 | 4 | 3 | 4 | 3 | 3 | **3,3** |
| | | 544 | 2 | 2 | 3 | 3 | 3 | **2,4** |
| | | **Average/lobe** | **3,3** | **3,3** | **3,5** | **3,5** | **3,0** | **3,3** |

**Supplementary Table 4. Hematological and biochemical parameters over time.** Values are presented as median and interquartile range (IQR). Statistical comparisons were performed using two-way ANOVA. P-values for group effects, time effects, and group–time interactions are reported in the first row for each parameter. Statistically significant p-values are shown in bold. 1. WBC, white blood cell count; 2. GGT, gamma-glutamyl transferase; 3. ALT, alanine aminotransferase; 4. ALP, alkaline phosphatase; 5. PT, prothrombin time; 6. INR, international normalized ratio.

| | Treatment | | | | | | | p-value | | |
|---|---|---|---|---|---|---|---|---|---|---|
| Parameter/ Time point | **PBS** | **PBS + VAP** | **PAO1 + PBS** | **PAO1 + CV8** | **PAO1 + CV8_VAP** | **PAO1 + CV8 (Nebulized)** | **PAO1 + CV8_VAP (Nebulized)** | **Group p-value** | **Time p-value** | **Interaction p-value** |
| WBC ($10^9$/L)1 | | | | | | | | | | |
| 0 hrs | 18.1 [18 - 18.3] | 15.7 [13.6 - 17.7] | 15.4 [13 - 35.2] | 16 [13 - 18.4] | 15.2 [11.8 - 15.6] | 7.7 [7.5 - 7.9] | 11.2 [10.5 - 18.8] | **0.006** | 0.41 | 0.30 |
| 12 hrs | 17.6 [17.4 - 17.9] | 13.3 [13.3 - 13.3] | 18.4 [16.5 - 23.3] | 14.1 [11.9 - 15.4] | 16.2 [11 - 17] | 11 [5.8 - 16.2] | 11 [7.7 - 14] | | | |
| 24 hrs | 20.4 [18.3 - 22.5] | 11.3 [10.9 - 11.8] | 12.9 [11.4 - 14.4] | 11.1 [8.5 - 12.5] | 11.9 [11.1 - 14] | 10.5 [7.5 - 13.6] | 12.1 [10.9 - 15.2] | | | |
| 36 hrs | 21.8 [16.8 - 26.8] | 11.1 [10.9 - 11.3] | 14.3 [10.5 - 15.6] | 11.5 [11.2 - 14.3] | 15.1 [12.7 - 16.7] | 11.8 [11.1 - 12.5] | 13.2 [12.3 - 25.2] | | | |
| Creatinine (mg/dL) | | | | | | | | | | |
| 0 hrs | 1.2 [1.1 - 1.3] | 1 [1 - 1] | 0.9 [0.8 - 1.1] | 0.9 [0.9 - 1] | 0.9 [0.9 - 1] | 0.7 [0.7 - 0.8] | 0.9 [0.8 - 1] | 0.51 | **0.009** | 0.31 |
| 12 hrs | 1.1 [1 - 1.1] | 0.8 [0.7 - 0.9] | 0.9 [0.8 - 1.1] | 1.1 [0.9 - 1.7] | 0.9 [0.9 - 1] | 0.8 [0.7 - 0.8] | 0.9 [0.7 - 1] | | | |
| 24 hrs | 1.2 [1 - 1.3] | 1.1 [1.1 - 1.1] | 1 [0.8 - 1.1] | 1 [0.9 - 2.8] | 1 [0.9 - 1.1] | 0.9 [0.7 - 1] | 1 [0.9 - 1.4] | | | |
| 36 hrs | 1.2 [1.1 - 1.3] | 1.2 [1.2 - 1.2] | 0.9 [0.8 - 1.1] | 1.2 [1 - 1.8] | 1 [0.9 - 1.1] | 1.6 [0.7 - 2.4] | 1.2 [0.9 - 2.3] | | | |
| Platelets (109/L) | | | | | | | | | | |
| 0 hrs | 440 [340 - 540] | 339.5 [235 - 444] | 535 [443.5 - 620] | 480 [366.5 - 531.5] | 375 [356.5 - 562] | 391 [357 - 425] | 404.5 [328.3 - 441] | 0.65 | **<0.001** | 0.27 |
| 12 hrs | 397.5 [341 - 454] | 274 [171 - 377] | 370 [293 - 434.5] | 292.5 [265 - 374] | 334 [281.5 - 455] | 344.5 [306 - 383] | 300.5 [237.5 - 421.3] | | | |
| 24 hrs | 380.5 [340 - 421] | 229 [174 - 284] | 360 [278 - 371] | 286 [265 - 326] | 308 [252.5 - 321.5] | 301.5 [273 - 330] | 309.5 [239.5 - 384] | | | |
| 36 hrs | 327.5 [298 - 357] | 228 [150 - 306] | 332 [240.5 - 346] | 292.5 [237.8 - 344.3] | 267 [234 - 300] | 257 [251 - 263] | 309 [280.3 - 364.8] | | | |

| | | | | | | | | | | |
|---|---|---|---|---|---|---|---|---|---|---|
| ALT (IU/L)2 | | | | | | | | | | |
| 0 hrs | 31.5 [30 - 33] | 28 [26 - 30] | 32 [26.5 - 36] | 33 [31 - 42] | 31 [22 - 35.5] | 27.5 [14 - 41] | 30.5 [17.3 - 34.8] | 0.57 | 0.11 | 0.44 |
| 12 hrs | 25 [25 - 25] | 23.5 [23 - 24] | 33 [30 - 34.5] | 34 [28.5 - 40] | 28 [21.5 - 33] | 29.5 [15 - 44] | 29 [16.3 - 31.3] | | | |
| 24 hrs | 26 [24 - 28] | 28.5 [24 - 33] | 33 [28 - 34.5] | 34 [28.5 - 42.5] | 28.5 [24 - 32.3] | 31 [16 - 46] | 22.5 [16 - 29.8] | | | |
| 36 hrs | 27 [23 - 31] | 29.5 [25 - 34] | 35 [27.5 - 39.5] | 33.5 [30.3 - 43.5] | 30 [23 - 31.5] | 32.5 [19 - 46] | 30 [18.5 - 34] | | | |
| GGT (IU/L)3 | | | | | | | | | | |
| 0 hrs | 30.5 [30 - 31] | 45.5 [24 - 67] | 41 [38.5 - 66.5] | 60 [35 - 101.5] | 47 [27 - 51.5] | 65 [59 - 71] | 36 [20.8 - 44.5] | 0.72 | 0.07 | **0.009** |
| 12 hrs | 78 [78 - 78] | 29.5 [24 - 35] | 61 [27 - 68] | 41 [32 - 58] | 32 [24 - 37.5] | 43.5 [22 - 65] | 33 [23.5 - 48.5] | | | |
| 24 hrs | 63 [35 - 91] | 37 [25 - 49] | 41 [25 - 45] | 31 [27 - 56.5] | 28 [19.8 - 32.5] | 44.5 [31 - 58] | 50.5 [33.5 - 82.5] | | | |
| 36 hrs | 54.5 [32 - 77] | 28 [22 - 34] | 28 [19.5 - 38.5] | 35.5 [26.3 - 56] | 30 [21 - 62] | 34 [29 - 39] | 29.5 [20.3 - 39.5] | | | |
| ALP (IU/L)4 | | | | | | | | | | |
| 0 hrs | 121 [112 - 130] | 214 [181 - 247] | 171 [121.5 - 207.5] | 190 [153.5 - 200.5] | 155 [109 - 251.5] | 70 [61 - 79] | 120 [96.8 - 126] | **0.04** | **<0.001** | **<0.001** |
| 12 hrs | 331 [331 - 331] | 176 [164 - 188] | 177 [112.5 - 185] | 179 [162.5 - 191.5] | 143 [92.5 - 238] | 65 [57 - 73] | 121 [83.3 - 137] | | | |
| 24 hrs | 195.5 [117 - 274] | 204.5 [166 - 243] | 151 [104.5 - 187] | 166 [155.5 - 184.5] | 174 [104.8 - 223.8] | 61.5 [53 - 70] | 106 [76 - 123.3] | | | |
| 36 hrs | 156.5 [102 - 211] | 192 [161 - 223] | 123 [95 - 166] | 143.5 [130.8 - 162.3] | 125 [81 - 193.5] | 58.5 [50 - 67] | 100.5 [68.5 - 151.3] | | | |
| PT (sec)5 | | | | | | | | | | |
| 0 hrs | 11.5 [10.9 - 12.1] | 11.4 [11.2 - 11.6] | 11.6 [10.5 - 11.9] | 11.7 [10.9 - 11.9] | 11.2 [10.4 - 12] | 11.1 [10.8 - 11.4] | 11.3 [11.1 - 11.4] | 0.12 | **<0.001** | 0.050 |
| 24 hrs | 11.7 [11.3 - 12.1] | 12.9 [12.8 - 12.9] | 13.1 [11.8 - 13.6] | 12.6 [12.5 - 12.8] | 12 [11.5 - 12.7] | 11.8 [11.2 - 12.3] | 11.3 [10.9 - 11.5] | | | |
| 36 hrs | 11.4 [11 - 11.7] | 12.6 [12.3 - 12.8] | 12.2 [12 - 13] | 11.9 [11.7 - 12.6] | 11.7 [11.5 - 12] | 11.7 [11.2 - 12.2] | 11.1 [10.7 - 11.7] | | | |
| INR6 | | | | | | | | | | |
| 0 hrs | 1 [0.9 - 1] | 0.9 [0.9 - 0.9] | 0.9 [0.9 - 1] | 1 [0.9 - 1] | 0.9 [0.8 - 1] | 1 [1 - 1] | 1 [1 - 1] | 0.95 | **<0.001** | **0.020** |
| 24 hrs | 1 [1 - 1] | 1 [1 - 1.1] | 1.1 [1 - 1.1] | 1.1 [1 - 1.1] | 1 [1 - 1.1] | 1 [1 - 1.1] | 1 [1 - 1] | | | |
| 36 hrs | 1 [0.9 - 1] | 1 [1 - 1.1] | 1 [1 - 1.1] | 1 [0.9 - 1.1] | 1 [1 - 1] | 1 [1 - 1.1] | 1 [0.9 - 1] | | | |

**Supplementary Table 5. Ventilator setting and adverse events during nebulization.**

| | PAO1 + CV8<br>N=2 | PAO1 + CV8_VAP<br>N=4 | p-value |
|---|---|---|---|
| Adverse events | | | |
| **Coughing (%)** | 1 (50%) | 0 (0%) | 0.333 |
| **Bronchospam** | 0 (0%) | 0 (0%) | >0.99 |
| **Oxygen desaturation** | 0 (0%) | 0 (0%) | >0.99 |
| **Mucus production** | 0 (0%) | 1 (25%) | >0.99 |
| **Discontinuation** | 0 (0%) | 1 (25%) | >0.99 |
| **Other adverse events** | 0 (0%) | 1 (25%) | >0.99 |
| Pre- and Post-arterial blood gases | | | |
| **Pre-arterial $PaO_2/FiO_2$** | 503.8 [474.8 – 532.8] | 405.9 [303.9 – 450.4] | 0.133 |
| **Post-arterial $PaO_2/FiO_2$** | 459.6 [444.3 – 475.0] | 373.3 [271.9 – 437.1] | 0.267 |
| **Pre-pH** | 7.6 [7.5 - 7.6] | 7.5 [7.4 - 7.5] | 0.133 |
| **Post-pH** | 7.5 [7.5 - 7.5] | 7.5 [7.4 - 7.5] | 0.533 |
| **Pre-arterial saturation** | 100.0 [99.9 – 100.0] | 99.7 [99.6 –99.9] | 0.208 |
| **Post-arterial saturation** | 99.8 [99.8 –99.8] | 99.7 [99.5 –99.7] | 0.067 |

#### SUPPLEMENTARY FIGURES

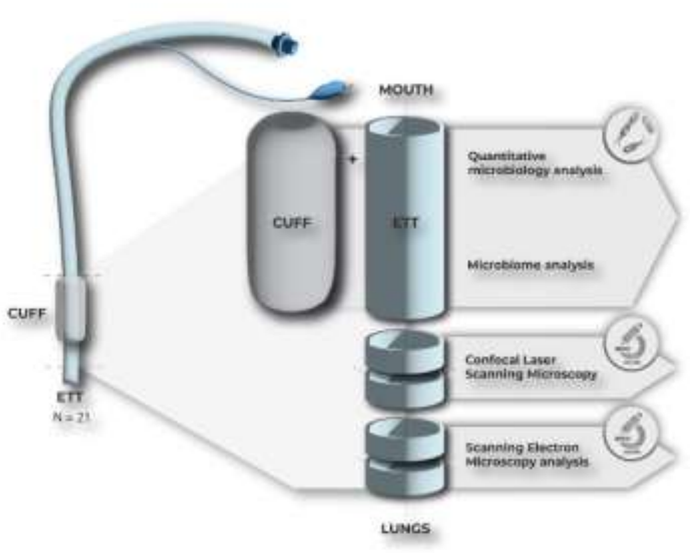


**Supplementary Figure 1. Slicing and sampling of the ETT and cuffs.** ETTs from mechanically ventilated pigs with VAP were obtained (N=21). All 3-cm sections and the corresponding cuffs were used for independent quantitative microbiological analysis (N = 21): PAO1 + PBS, n= 5 (all intralobe); PAO1 + CV8, n = 7 (intralobe, n = 5, nebulized, n = 2); PAO1 + CV8_VAP, n = 9 (intralobe n = 5, nebulized n = 4). The 3-cm sections were also used for the microbiome analysis (N = 20, excluded one ETT sample from PAO1+CV8_VAP because of low quality DNA sample). All 0.5-cm cross sections were analyzed by SEM and were observed by CLSM.

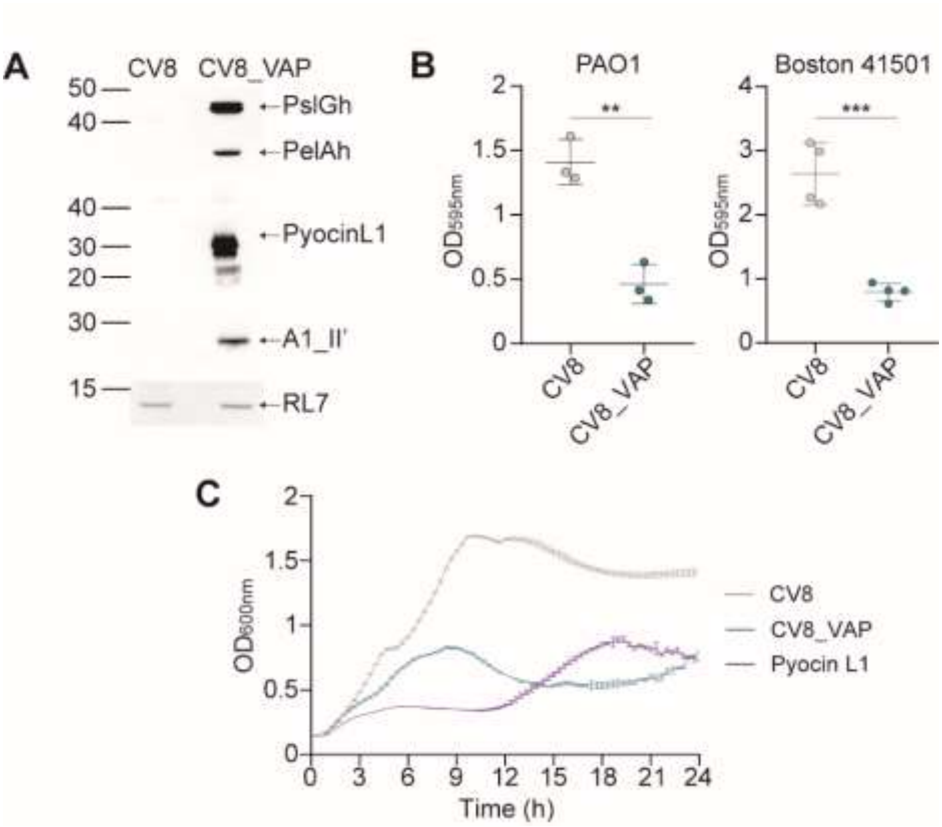


**Figure S2.** ***In vitro*** **characterization of the CV8_VAP strain. (A)** Western blot showing the expression of the payloads in the cell lysate of the CV8_VAP strain compared to the control strain CV8. The ribosomal protein RL7 was used as a loading control. Membranes were blocked with 5% skim milk in TBS-Tween and incubated with either a monoclonal anti-FLAG antibody (Sigma, F1804) to detect PelAh (28.2 KDa) and PslGh (48.7 KDa), or custom-made anti-A1-II' (24.7 KDa) and anti-pyocin L1 (28.3 KDa) polyclonal primary antibodies (Proteogenix). **(B)** Crystal violet assay demonstrating the anti-biofilm activity of the CV8_VAP supernatant against *P. aeruginosa* PAO1 (left) and Boston 41501 (right) strains. Data are presented as mean ± SEM (n = 3). Statistical significance was determined using Student's t-test: **p < 0.01; ***p < 0.001. **(C)** Growth curve of *P. aeruginosa* PAO1measured by absorbance at $OD_{600}$ when incubated with the supernatant of CV8 (grey), CV8_VAP (blue), or recombinant pyocin L1 protein (47.5 nM, purple).

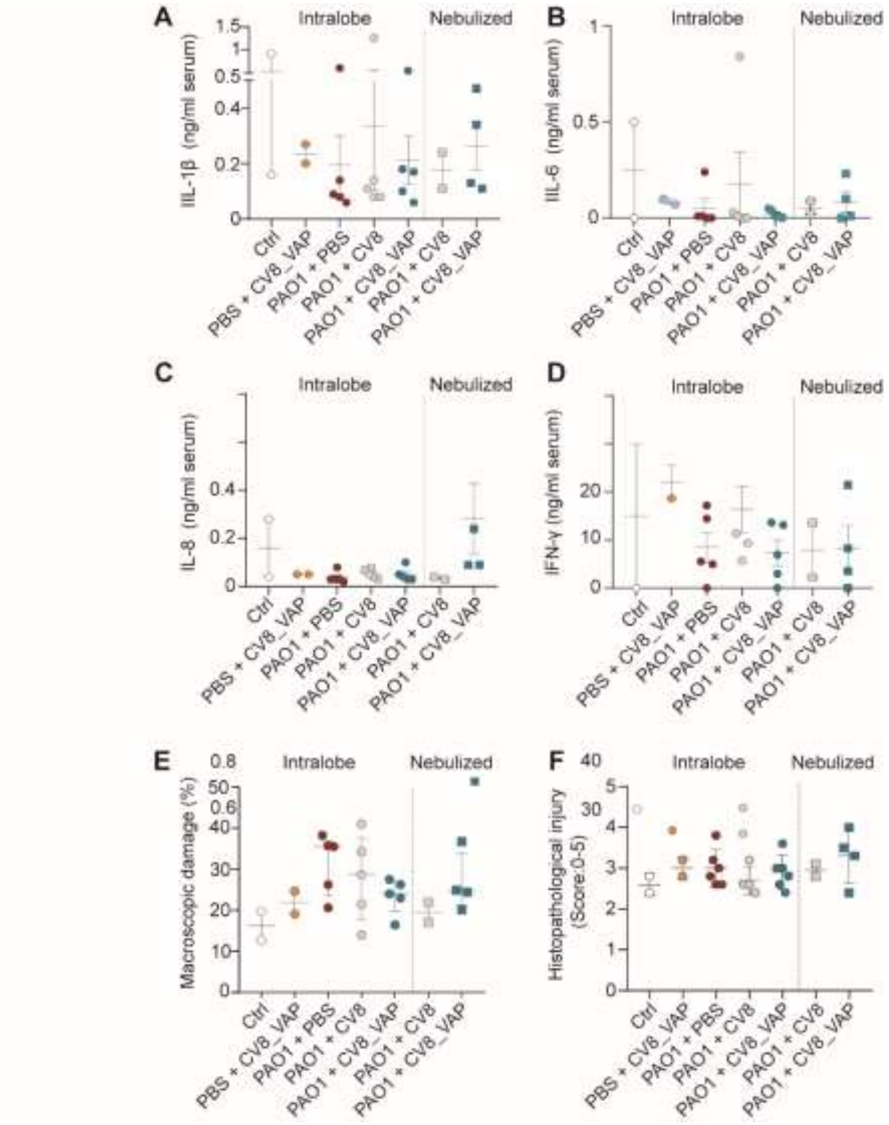


**Supplementary Figure 3. Impact of the VAP Platform on *P. aeruginosa*-Induced lung inflammation and pulmonary injury. (A-D)** Inflammatory proteins detected in serum samples (ng/mL) at the time of autopsy (36h). Macroscopic (**E**) and histopathological (F) evaluation of lung tissue. The histopathology raw data are provided in Supplementary Table 3.

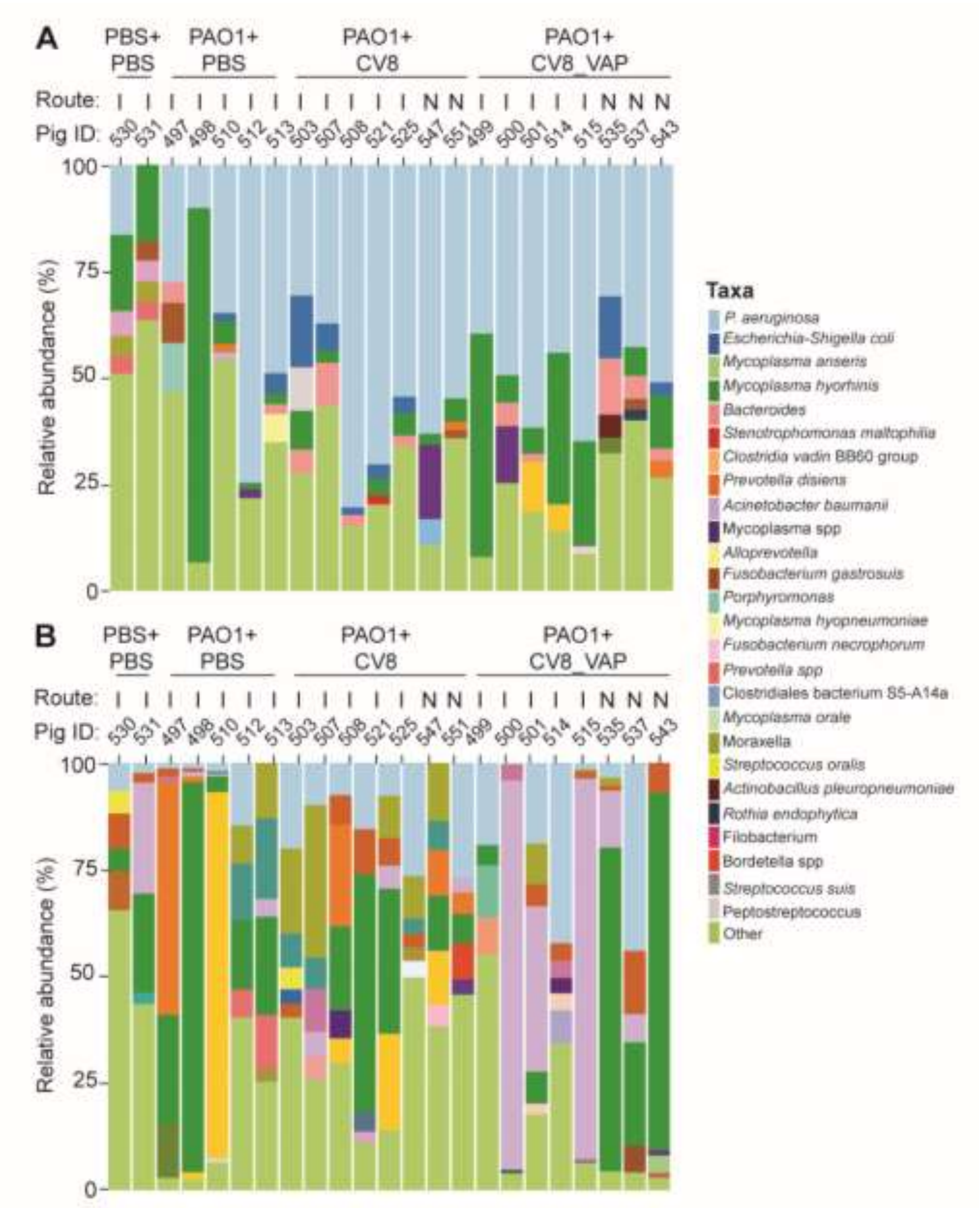


**Supplementary Figure 4. Relative abundance of top taxa identified in ETTs (A) and lung (B).** Taxa are shown at the highest taxonomic resolution possible (genus and species). In the top, each individual animal and its treatment group: PAO1 + PBS (n = 5; intralobe), PAO1 + CV8 (n = 7; intralobe, n = 5, nebulized, n = 2), and PAO1 + VAP (n = 8; intralobe, n = 5, nebulized, n = 3). Left Y-axes show relative abundance. Intralobe (I), Nebulized (N).
